# Polarity-JaM: An image analysis toolbox for cell polarity, junction and morphology quantification

**DOI:** 10.1101/2024.01.24.577027

**Authors:** Wolfgang Giese, Jan Philipp Albrecht, Olya Oppenheim, Emir Bora Akmeriç, Julia Kraxner, Deborah Schmidt, Kyle Harrington, Holger Gerhardt

**Affiliations:** Max Delbrück Center for Molecular Medicine in the Helmholtz Association, Berlin, Germany; DZHK (German Center for Cardiovascular Research), Berlin, Germany; HELMHOLTZ IMAGING, Max Delbrück Center for Molecular Medicine in the Helmholtz Association, Berlin, Germany; Humboldt-Universität zu Berlin, Faculty of Mathematics and Natural Sciences, Berlin, Germany; Charité - Universitätsmedizin Berlin, Berlin, Germany; Chan Zuckerberg Institute for Advanced Biological Imaging, Redwood City, CA, USA; Berlin Institute of Health, Berlin, Germany

**Keywords:** Cell polarity, Image analysis, Circular statistics, Endothelial cells, Software, Workflow

## Abstract

Cell polarity involves the asymmetric distribution of cellular components such as signaling molecules and organelles within a cell, asymmetries of a cell”s shape as well as contacts with neighbouring cells. Gradients and mechanical forces often act as global cues that bias cell polarity and orientation, and polarity is coordinated by communication between adjacent cells.

Advances in fluorescence microscopy combined with deep learning algorithms for image segmentation open up a wealth of possibilities to understand cell polarity behaviour in health and disease. We have therefore developed the open-source package Polarity-JaM, which offers versatile methods for performing reproducible exploratory image analysis. Multi-channel single cell segmentation is performed using a flexible and userfriendly interface to state-of-the-art deep learning algorithms. Interpretable single-cell features are automatically extracted, including cell and organelle orientation, cell-cell contact morphology, signaling molecule gradients, as well as collective orientation, tissue-wide size and shape variation. Circular statistics of cell polarity, including polarity indices, confidence intervals, and circular correlation analysis, can be computed using our web application. We have developed data graphs for comprehensive visualisation of key statistical measures and suggest the use of an adapted polarity index when the expected polarisation direction or the direction of a global cue is known *a priori*.

The focus of our analysis is on fluorescence image data from endothelial cells (ECs) and their polarisation behaviour. ECs line the inside of blood vessels and are essential for vessel formation and repair, as well as for various cardiovascular diseases, cancer, and inflammation. However, the general architecture of the software will allow it to be applied to other cell types and image modalities. The package is built in in Python, allowing researchers to seamlessly integrate Polarity-JaM into their image and data analysis workflows, see https://polarityjam. readthedocs.io. In addition, a web application for statistical analysis, available at www.polarityjam.com, and a Napari plugin are available, each with a graphical user interface to facilitate exploratory analysis.

## Introduction

Cellular polarity is important in many biological phenomena spanning from developmental processes to wound healing and angiogenesis. Cell migration, cell division, and morphogenesis depend on prior polarisation and breaking of spatial symmetry. Spatial reorganisation of the plasma membrane, cytoskeleton, cell-cell junctions, or organelles is required to establish an axis of polarity with a distinct direction, meaning “front and back”, to guide directed processes (1). In these processes, cells have to adapt and react according to multiple and often conflicting cues from their environment (2).

Fluorescence microscopy has become an invaluable tool for producing high-resolution, high-content images of *in vitro* systems as well as *in vivo* tissues. These images can be obtained at subcellular resolution of less than one micron, with multiple fluorescence channels acquired in parallel, often on multiple planes, allowing for detailed quantification of cellular polarity and asymmetries. At the same time, deep learning segmentation algorithms have developed at a staggering pace over the past few years, enabling the segmentation of individual cells and organelles with near-human accuracy (3– 8). This opens up a wealth of unprecedented possibilities, but also creates new challenges for comprehensible and interpretable image data analysis workflows that fully exploit these new potentials.

Each cell has a unique shape, a particular spatial distribution of organelles, and contains different absolute amounts (measured by intensity) and distributions (gradients) of protein species. In addition, junctional cell-cell contacts exist with their own morphological phenotypes. Taken together, this image-based information provides a snapshot of a cell”s state, which we aim to turn into quantitative and comparable features. We demonstrate our investigations on image data from endothelial cells (ECs). ECs line the inside of blood vessels and play a crucial role in organ function and health of the whole organism. Migration of these cells is important for vessel formation and repair, but can also be involved in disease processes in the cardiovascular system, cancer, or inflammation. ECs experience shear stress when blood flow passes through blood vessels, causing alterations at the collective and single cell level, including changes in EC polarity, alignment, shape, size, junctions, and gene expression. We describe some of the measurements and their biological correlates, see Fig. 1. The location or distribution of the organelle in the cell may be quantified in relation to the nucleus or the cell centre. For example, the position of nucleus and Golgi apparatus are used to compute orientation vectors for each cell, see Fig. 1 A,B. Nuclei-Golgi polarity has been positively correlated with directed motility (9) and cell orientation in tissue (10) in several cell types, including endothelial and epithelial cells. EC polarity can be induced by multiple signaling cues, including shear stress and VEGFA, which can be in competition with each other (11). Various EC shear stress sensors have been identified that induce EC polarisation, including Piezo1, plexinD1, focal adhesions (FA), the VEGFR2 / PECAM1 / VE-cadherin complex, or caveolae (12). As a result, ECs adapt their shape and orientation to shear forces (10, 13). Cell shape is often measured using circularity, shape index, or length-to-width ratio (LWR). The LWR has been positively correlated with VEGFA treatment in all EC types (14), the collective signature also differs across organs and microenvironments (15). Collective cellular orientation has also been correlated with both shear flow types, pulsatile and laminar, which have been shown to organise collective EC phenotypes. Pulsatile flow does not organise the direction of orientation as strongly as laminar flow (16, 17).

**Fig. 1.**
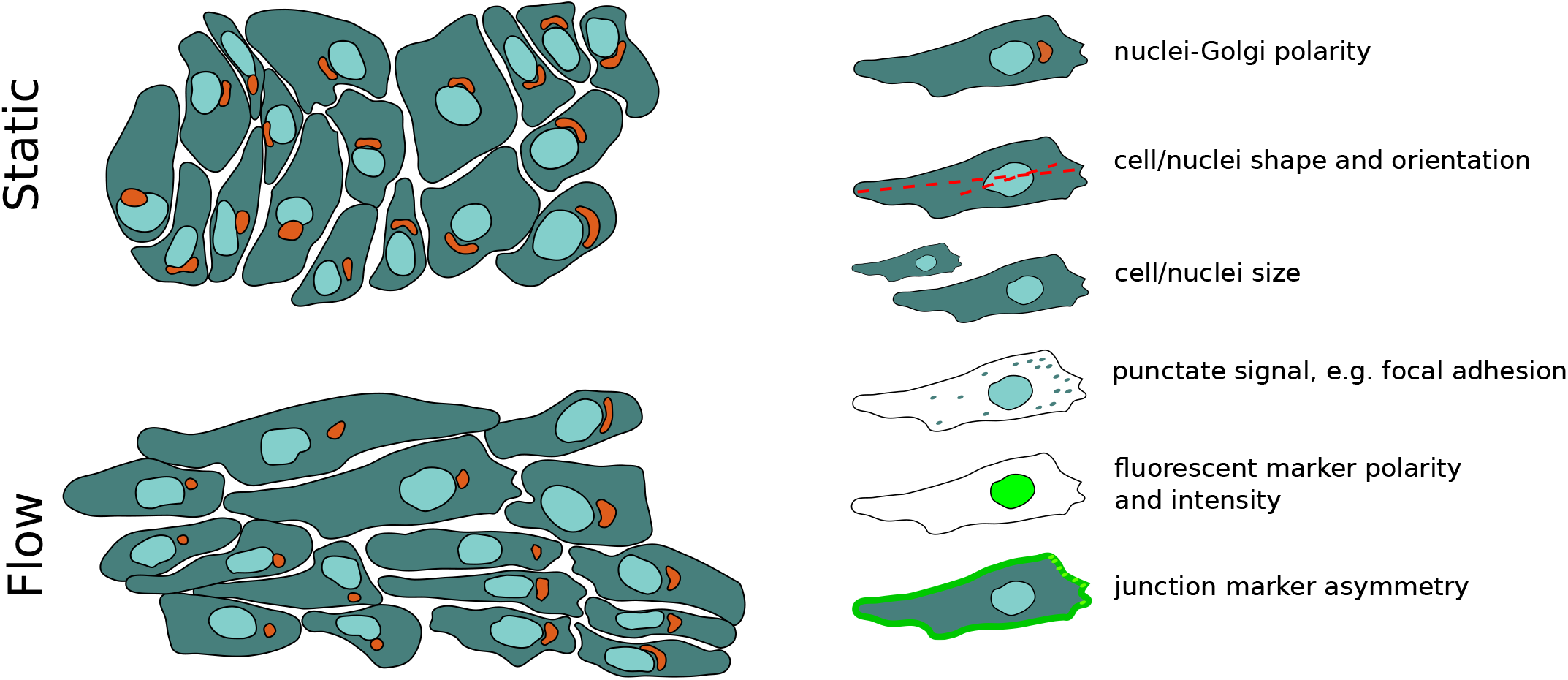
**A**: Endothelial cells under static conditions have a random polarity, while endothelial cells exposed to flow (the direction is indicated by the black arrow) are polarised against flow (indicated here by the relative nuclei-Golgi polarity) and elongate. **B**: Examples of image-based cellular polarity, morphology, and cell-cell junction readouts in Polarity-JaM.

A major challenge in the creation of every image analysis pipeline is identifying and measuring informative features. This search has a large iterative component and is based on precise and accurate measurement of the relevant microscopy data (18). Several studies have proposed new meaningful measures such as Quantify Polarity (19), Junction Mapper (20) and Griottes (21) or reviewed existing ones (18). However, it is not yet possible to integrate these different aspects into a single pipeline and perform a multivariate analysis. These requirements and constraints motivated the development of the Polarity-JaM package, which is built to streamline the process of exploratory image analysis; this is accomplished by providing the end user with functionality that includes a wide range of features, explanatory metadata, clear and concise documentation, and high testing coverage. Proper meta-data and testing mean that parameters can be tuned safely and systematically, and the functionality of the three main components of the package can be -reasonably-extended to new types of analysis. All relevant analysis can be performed with our package, making installation and usage straightforward. For relevant terminology in our article, we provide a glossary at the end of the supplement, see Table S14.

To visualise and explore the wide range of cellular polarity, morphology, and junction features, we have developed an R-shiny application. In particular, our toolbox focusses on polarity data, which typically requires bespoke statistical tools for circular data that are not available in standard toolboxes. Circular statistics and correlations can be computed and visualised in scatter plots. We make use of phenotypic descriptors that provide the user with bonafide summary statistics that can be correlated with biological processes. For example, flow-mediated morphogenesis in EC collectives, which affects descriptors of cell elongation, cell shape, and nuclei orientation, as well as nuclei-Golgi polarity and changes in junctional morphology, can be directly correlated using circular statistics in our Web App. A central aim of our package is to aggregate diverse biologically descriptive statistics under the auspices of a single workflow that can measure multiple aspects of endothelial biology. In our Methods and Results section, relevant statistics are described and their combined use demonstrated.

In this article, we present Polarity-JaM, an open-source software suite for measuring cell features to describe collective behaviour in microscopic images. The engineering of features is a non-trivial task and requires biological expert knowledge as well as deep understanding of the data and problem, which is explicitly addressed in our proposed pipeline. We will discuss several examples typical for EC flow assays, including quantification of polarity in subcellular organelle arrangement, orientation of cell and nuclei shape, subcellular signal asymmetries based on intensity measurements, subcellular localisation of signaling processes, and cell-cell junction morphology. In parallel, we will introduce informative statistical plots that allow to display all relevant information, including a circular histogram, the polarity index, confidence intervals and circular correlations - many of these useful measures are often omitted or rarely used in the quantification of collective cell systems. In summary, we propose a holistic image analysis workflow that is accessible to the end user in bench science.

## Results

### Asymmetries in subcellular localisation of organelles

Cell polarisation is a dynamic process that involves the reorganisation of various cellular components, including the cytoskeleton, intracellular signaling molecules, organelles, and the cell membrane. The nucleus is the most prominent organelle in eukaryotic cells and is constantly exposed to intrinsic and extrinsic mechanical forces that trigger dynamic changes in nuclei morphology and position (22). The centrosome is outside the nucleus, also referred to as the microtubule organisation centre (MTOC) and, unlike its name, is rarely located at the geometric centre of the cell (23, 24). The Golgi apparatus is involved in the establishment of cell polarity by sorting and packaging proteins into distinct membranebound vesicles that are then trafficked to different parts of the cell (25). In most animal cells, the Golgi apparatus is one complex, and its position is frequently used as a read-out for cell polarity and indicator of migration direction. Note that there are several other organelles in other cell types that could be used as readouts, which are not discussed here. In summary, organelles are positioned around the nucleus, and their positions are crucial to many directed processes, such as cell division, directed migration, asymmetric adhesion, or heterogeneous cell-cell contact. Their positioning may have passive or active effects on crucial mechanisms, including Rho GTPase signaling, but also Map Kinase signalling (26, 27). Sprouting angiogenesis and vascular remodelling are based on EC polarity and directed migration (28, 29). A common readout in microscopic images of ECs *in vivo* and *in vitro* are relative nuclei-Golgi locations. In response to flow-induced shear stress, the Golgi is located upstream of the nucleus against the flow direction (10). The nuclei-Golgi polarity is often used as a proxy for the migration direction in static images.

We will exemplify our approach using nuclei-Golgi positioning and nucleus displacement as read-out; see Fig. 2. We applied shear stress levels of 6 dyne/cm^2^ and 20 dyne/cm^2^ for 4 h under different media conditions, to induce robust collective polarisation of the EC monolayers (see Fig. 2). Microscopic images contained a junction channel, a nuclei channel, and Golgi staining. An example image is shown in Fig. S1A and Fig. S2. Using the Cellpose (4) algorithm, we obtained segmentations for cells and nuclei. Golgi segmentations were obtained by applying Otsu thresholding directly to the Golgi stained channel and superimposing the resulting mask with the cell segmentation from Cellpose. The results are shown in Fig. S1 B. The nuclei-Golgi vectors were automatically calculated for each cell, see Fig. 2 A. In tandem, we also obtained vectors from the cell centres to the nuclei centres, which provide a measure of the direction of nuclei displacement.

**Fig. 2.**
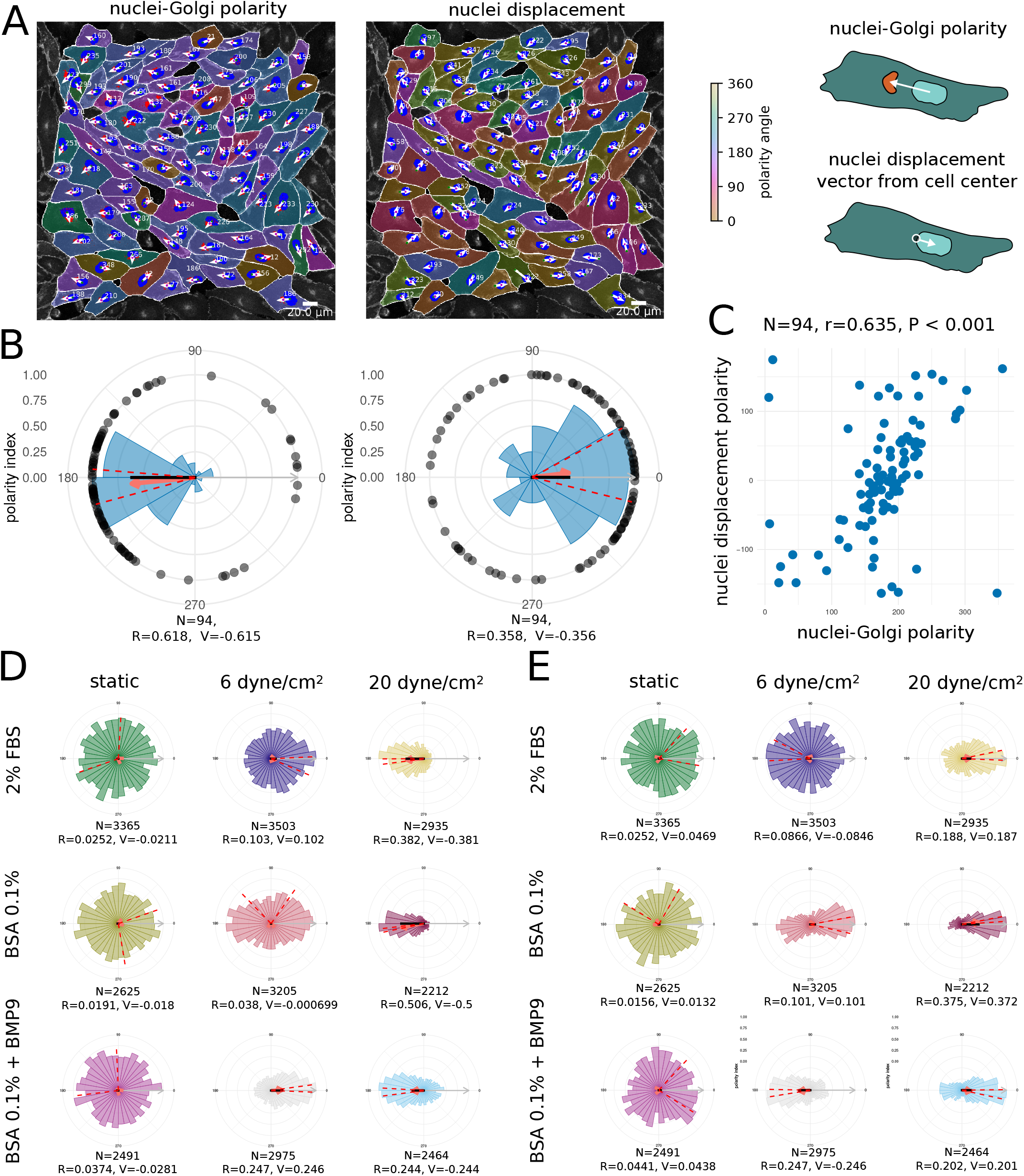
Asymmetric localisation of organelles as a measure for cell polarity. **A**: Human umbilical vein endothelial cells (HUVECs) in culture exposed to 20 dyne/cm^2^ shear stress at 4 h. Nuclei-Golgi polarity [left] and displacement of the nuclei within each cell with respect to its centre [right]. Orientation is indicated by a cyclic colour scheme and white arrows from the centre of the nuclei to the centre of the Golgi (nucleus-Golgi polarity) or from the centre of each cell to the nucleus (nucleus-displacement polarity). The flow direction is always from left to right. Scale bar 20 µm . **B**: Circular histograms of the distribution of cell orientations of a single image; the red arrow indicates the mean direction of the cell collective and its length is the polarity index. The red dashed lines indicate the 95% confidence intervals. Black dots indicate single-cell measurements. The grey arrow indicates the polar vector that points in the direction of flow, and the length of the black bar indicates the signed polarity index (V). **C**: Circular correlation of nuclei-Golgi polarity and nuclei displacement polarity. **D**: Ensemble plot of nuclei-Golgi polarity generated with the Polarity-JaM app for different flow conditions, each containing several hundreds of cells from *N*_*r*_ *≥* 3 biological replicates. **E**: Ensemble plot of the nuclei displacement orientation polarity generated with the Polarity-JaM app for different flow conditions, each containing several hundreds of cells from *N*_*r*_ *≥* 3 biological replicates.

The collective strength of polarisation is commonly measured using the polarity index (30), which is calculated as the resultant vector of all orientation vectors from each single cell. Mathematically we obtain a vector for every single cell, which in our example points from the nucleus to the Golgi:

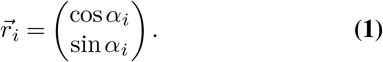

The average of the individual vectors is used to calculate the resultant vector from

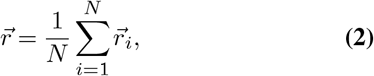

where *N* is the number of cells. The length of this vector, computed from 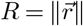 is called the polarity index (PI) and its direction is the mean of the distribution. The value of the polarity index varies between 0 and 1 and indicates how much the distribution is concentrated around the mean direction. A polarity index close to 1 implies that the data are concentrated around the mean direction, while a value close to 0 suggests that the data are randomly distributed or spread in several directions. In summary, the polarity index indicates the collective orientation strength of the cell layer or tissue. Note that the polarity index is closely related to the variance of the distribution by *S* = 1 *− R*. In Fig. 2 B,D,E the value of the polarity index is shown and indicates the strength of the collective flow response.

To complement this analysis, we introduce a signed polarity index, which is a modified version of the V-score and is calculated to obtain the V-test statistics (31). The signed polarity index assumes a known polar direction. For example, if we assume that flow is orientated from left to right and Ecs polarise against flow, we set the polar direction to an angle of *µ*_0_ = 0 in our reference system, see Fig. 2 B. The signed polarity index is computed from:

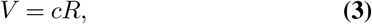

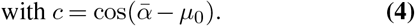

The signed polarity index varies between -1 and 1 and indicates the strength of polarisation with respect to an assumed polarisation direction. In our example, a value of -1 indicates that all cells are perfectly orientated against flow, while a value of 1 indicates that all cells are perfectly orientated with flow. For values in between, the distribution is more spread or diverges from the polar direction. Therefore, the signed polarity index measures both the deviation from the expected polarity direction and the spread of the distribution. We suggest a graphical design for the representation of the polarity index, the signed polarity index, the polar direction, confidence intervals, circular histogram, and single measurements, see supplementary Fig. S3.

The distribution of the polarity index (R) and signed polarity index (V) for *N*_*i*_ = 30 images per condition is shown in Fig. S4 A, B for the nuclei-Golgi polarity and in Fig. S4 C,D for nuclei displacement. We found that the nulcei-Golgi polarisation of the ECs under shear stress is highly correlated with the displacement of the nuclei and points in the opposite direction, Fig. 2 B,C. We found the same behaviour for a wide range of flow conditions; see Fig. 2 D,E and Fig. S4 E, but with different magnitudes depending on the media conditions and magnitude of shear stress. Our web application includes a number of different statistical tests, including the Rayleigh test, the V-test, Watson test, and Rao”s spacing test, see Supplementary Note 6. It should be noted that these tests are not appropriate for measurements of individual cells, given that the groups of single cells within a monolayer or tissue exhibit significant correlation. We therefore recommend the use of estimation statistics (32) to calculate effect sizes Fig. S4 of collective parameters such as the polarity index (R) and signed polarity index (V).

In (33) a systematic analysis has demonstrated that the V-test, which is based on the computation of the signed polarity index, is recommended over other tests if an expected direction is known *a priori*. Note that here we use the flow direction as the polar direction, whereas for the V-test the expected polarisation direction must be used. For the nuclei-Golgi polarity, for example, the expected polarisation direction points in the opposite direction to the flow direction, while for the nuclei displacement both the flow direction and the expected polarisation direction are the same. The signed polarity index in our data provides a more accurate or comparable distinction of all conditions with respect to control, in particular for 6 dyne/cm^2^ and 20 dyne/cm^2^. In conclusion, it is advantageous to use the signed polarity index when the expected direction of polarisation or the direction of an external signal is known in advance.

### Shape asymmetries

Cell morphology is an important aspect of cell biology as it provides insight into the structure and function of a cell. Cell morphology can be used to study the effects of different treatments on cells, the effects of genetic mutations, to understand how cells interact with their environment, and to identify potential therapeutic targets for disease. Furthermore, the cell shape is strongly connected to cell polarity, direction of cell movement, orientation of cell division, and organisation of the cytoskeleton. The shape of the cell also affects the ability of cells to interact with their environment and to respond to external signals. There are also different effects on the organisation of molecules, the protrusion can function as small pockets and reaction chambers, while signaling gradients are more easily stabilised along the long axis of the cell than the short axis (27, 34). Cell shape is also coupled with the migration direction; for example, in keratocyte-like motion, polarisation occurs along the short axis of the cell (35), while other cell types migrate along the long axis (36).

Recent studies have shown that monolayer orientation is modulated by both physical and chemical stimuli. Vion et al. (17) show that shear stress and VEGF-A act to modulate the cohesiveness of cell orientation in the endothelial monolayer. Physiochemical stimuli interact to modulate patterns of EC interaction across scales. As shear stress increases, endothelial cells polarise against flow, however, this phenotype is ablated with EC senescence. Furthermore, Deb et al.

(16) demonstrated that cell size is modulated by shear flow. Longer exposure to shear flow decreases cell area, while increasing cell aspect ratio across longer (approx. 48 h) time scales. The size of ECs has also been positively correlated with senescence (37). Laminar shear stress induces endothelial cell elongation (38). Alteration in substrate stiffness leads to changes in cell shape through mechanotransduction (39). Furthermore, cell shape can be an indicator of the mesoscale properties of tissue or cell monolayer and is often used as an order parameter in soft matter physics (40). In summary, cell shape and orientation are important readouts for directed cellular processes.

We applied our workflow to endothelial cells exposed to 6 dyne/cm^2^. The angle of the major axis of the cell or nucleus shape with the x axis serves as a readout for their orientation, see Fig. 3A,B. Here, any angle *α*_*i*_ is identified with its opposite *α*_*i*_ + 180, thus we do not distinguish the front and back of the cell (or nucleus). An angular histogram showing the angle distribution was then generated with our app, which in this case is mirrored on the x-axis. Note that angular data of cell shape and nuclei orientation angles are referred to as axial data, which means that all orientation angles *α*_*i*_, 1, *· · ·, N* take values between 0 and 180 degrees. These axial orientation data were converted to directional data by doubling all values *θ*_*i*_ = 2*α*_*i*_. The mean direction was calculated from 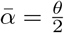, where 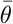 is the common circular mean of the directional data *θ*_*i*_. Similarly, the polarity index (R) was calculated as the length of the mean resulting vector of directional values *θ*_*i*_. Again, the polarity index varies between 0 and 1 and indicates how much the distribution is concentrated around the mean. A polarity index close to 1 implies that the data are concentrated around the mean direction, while a value close to 0 suggests that the data are evenly distributed or random. The V-score can be computed in the same fashion as for directed circular data (see Supplementary Note 6 for more details).

**Fig. 3.**
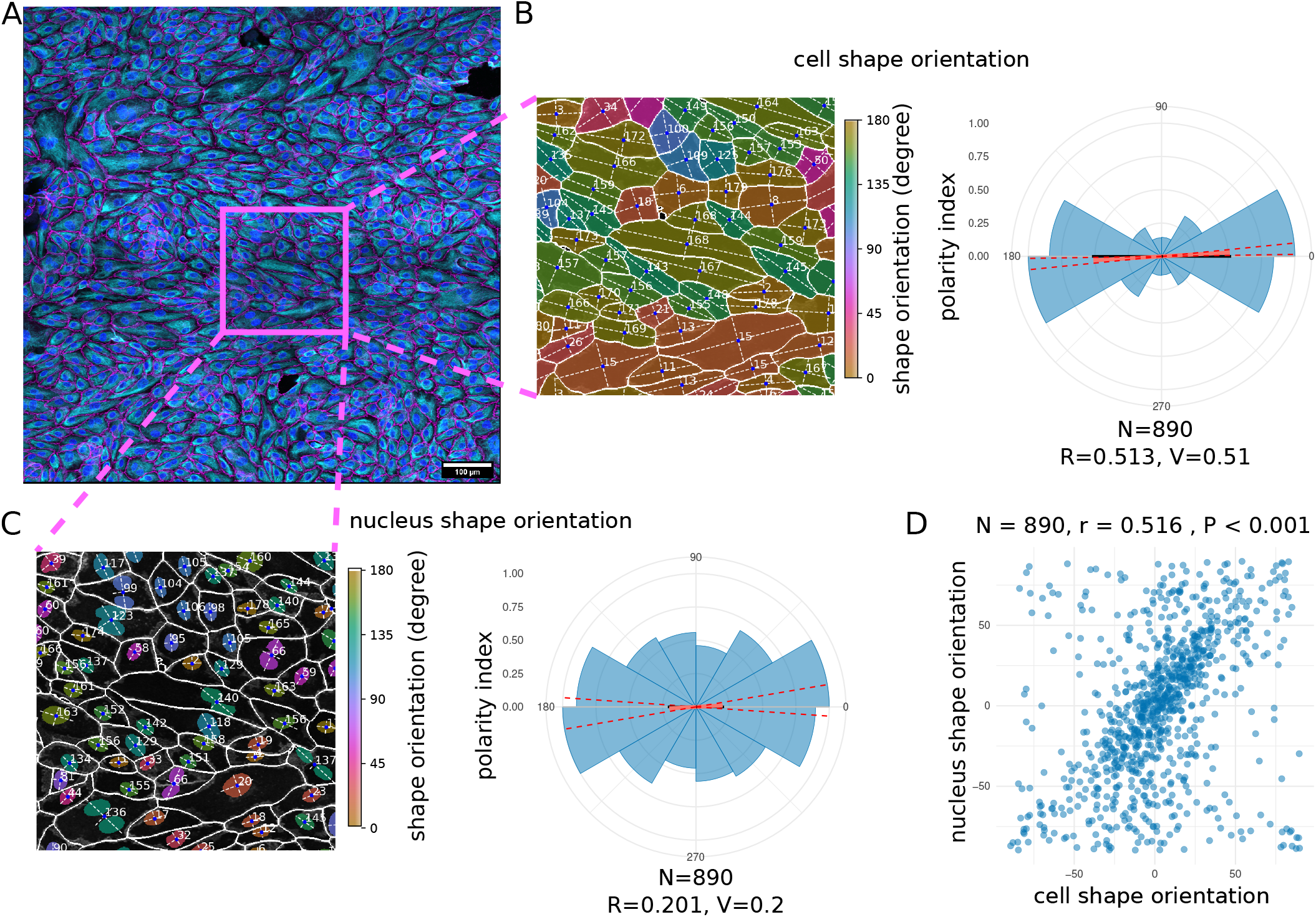
Asymmetry in cell and nuclei shape and their orientation. **A:** Example input image. **B**,**D:** The major and minor axes of the cell (B) and nuclei (D) shapes are indicated with white dashed lines. The angle of the major axis with the x-axis serves as a readout for the orientation of the cells (B) and nuclei (D), which is indicated with a circular colour scheme ranging from 0 to 180 degrees. **C:** Circular correlation between cell shape orientation and nuclei orientation.

Changes in intracellular state or interactions with neighbouring cells can also induce changes in the shape of the cell and its organelles, in particular the nucleus (22). We tested the coupling of cell shape orientation to nuclei orientation in ECs and found a significant correlation, see Fig. 3. Similarly, we found that cell elongation and nuclei elongation are coupled. This shows that nuclei morphology and orientation can serve as a proxy for shape elongation and orientation.

There are different commonly used measures of shape changes that we divide into two groups. The first group is determined by fitting an ellipse to the cell shape, identifying its major (longest) and minor (shortest) axes. However, this approach does not capture the complexity of protrusions such as filopodia or lamellipodia. The second group is computed from the relation of area and perimeter, such as, for example, circularity or shape index (40), see Materials and Methods for more details. In particular, the shape index has been popular in soft matter physics to characterise the mesoscale properties of tissues or cell monolayers as solid or fluid (41).

### Quantification of intracellular signalling gradients

Gradients or asymmetric distributions of signaling molecules are inherent in cell polarity. Often these asymmetries are decoded into rather small and subtle gradients that can be amplified by signaling feedback systems, such as the Rho GTPase system (42, 43). The establishment of these signaling gradients within single cells allows cell collectives to respond to their environment in a coordinated manner and is used to control cell migration, differentiation, and other cellular processes (44). Quantifying the direction and strength of intracellular signaling gradients between different experimental conditions is therefore crucial to gain insight into the underlying processes.

The notch signaling pathway is involved in the regulation of various genes responsible for angiogenesis (45) and has been shown to be a mechanosensor in adult arteries (46). Therefore, the effects of this pathway are of great interest. We present an automatic quantification approach of signaling gradients for each cell with the example of the NOTCH1 protein using our tool. We investigated both circular and linear features. First, the marker polarity is a novel circular feature, which can be described as the direction from the geometric cell centre to the weighted centre of marker intensity of the cell. Second, we computed the cue directional intensity ratio as a linear feature, which can loosely be described as the ratio between the mean intensities of the left-hand and right-hand cell-half of a cell perpendicular to a given cue direction. We provide a detailed description in the Materials and Methods section. The ratio takes values ranging from [-1, 1] where -1 indicates a strong asymmetry against a direction of the cue, 0 without visible effect, and +1 a strong asymmetry along the direction of the cue. An example image together with a visualisation of the cue directional intensity ratio and marker polarity is shown in Fig. 4 A (from left to right). A similar approach of using ratios on opposite sides of the cell was introduced by (47) to determine the magnitude and angle of polarity of a given cell.

**Fig. 4.**
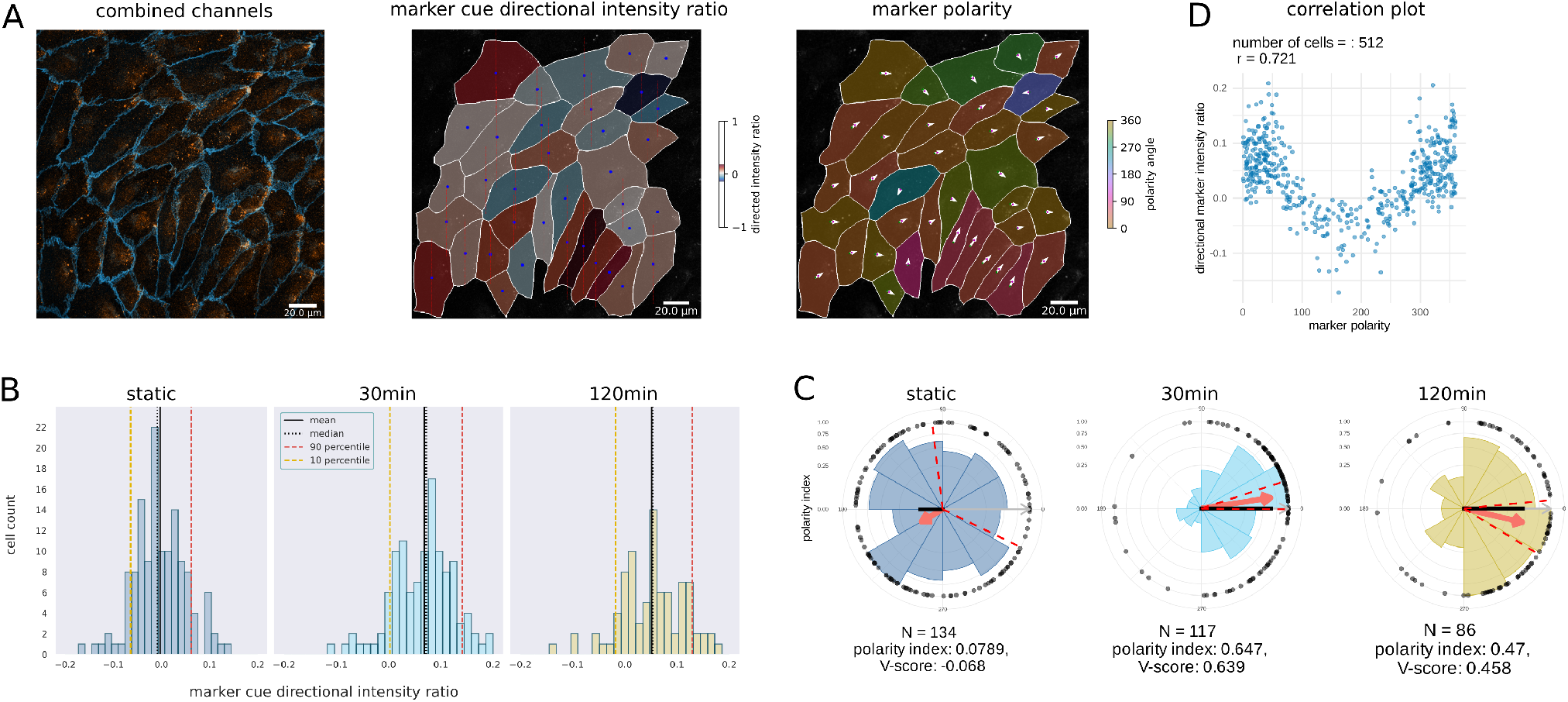
Notch signal signaling asymmetry in endothelial cell culture exposed to flow. **A:** Left shows the NOTCH1 signal (orange) combined with the junction channel (blue). Middle shows the NOTCH1 (here called marker) cue directional intensity ratio. The blue circle inside each cell marks the centroid of the cell, while the vertical dashed red lines are perpendicular to the flow direction and thus separate the cell to front and back. Right shows the marker polarity as a directional property. **B:** Shows the cue directional intensity ratio of the NOTCH1 signal with respect to the flow direction before [left], after 30 minutes [middle], and after 2h [right] of exposure to shear stress. Statistics mean +std from left to right: -0.0026+-0.0539, 0.0687+-0.0597, 0.0512+-0.0609. **C:** The circular histogram of the polarity of the NOTCH1 signal before applying shear stress [left], after 30 minutes of exposure [middle], and after 2 h of shear stress [right]. The red arrow points towards the collective mean direction of polarity, and its length represents the polarity index. Dashed lines indicate the 95% confidence intervals of the circular mean. Black dots indicate single-cell measurements. **D:** Correlation between marker polarity and directional intensity ratio.

To investigate the polarity of the Notch1 signal, we compared ECs in static conditions and exposed to shear stress for a 2 h time period. The images contained a junction and nucleus channel as well as a NOTCH1 staining. The junction and nucleus channel were used for segmentation to then quantify the intracellular NOTCH1 signaling gradient.

The result of the analysis is shown in Fig. 4 bottom row. We observed a shift in the marker cue directional intensity ratio when comparing static and shear stress conditions. We observed a rather small, but consistent change in mean, indicating a small gradient of the NOTCH1 signal, with a lower concentration on the flow facing side, Fig. 4 B. The results of the marker polarity analysis showed strong asymmetry effects, see Fig. 4 C, with large polarity index of 0.647 and 0.47 and V-score of 0.639 and 0.458 at 30 min and 120 min, respectively, while the mean was pointing along the direction of flow. This confirms the results from (46). A circular-linear correlation analysis between the polarity index and the directional intensity ratio of the cue revealed a correlation coefficient of 0.721, see Fig. 4 D, which implies a strong correlation between both measurements. Since signaling gradients in a single cell are rather small and sometimes noisy(48), ratio values are expected to be close to zero and show higher variance. However, the cue directional intensity ratio is a linear feature, easy to interpret, and can be compared between experimental setups, which is why we included it in our set of features. The marker polarity, on the contrary, is much more sensitive, but does not measure the magnitude of the gradients, but only their direction.

### Localised marker expression of KLF4

To complement our investigation of signal intensity gradients, we also characterised the localisation of image-based signal intensities, which indicate the localisation of specific processes. Localisation of cellular processes in biological cells is important because it allows for precise regulation of downstream processes, such as gene expression, protein synthesis, signaling and other cellular activities. For instance, it is important whether molecules are localised at the cell membrane, where they might get activated via phosphorylation, or if they fulfil other functions through anchoring to the membrane. Localisation to the nucleus is also important for a variety of cellular processes, including gene expression or RNA processing.

To quantify the signal in the different subcellular compartments, we compute the total amount and concentration of signal intensity, in the nucleus, the cytosol (without the nucleus) and the membrane nucleus, see Fig. 5 A. We demonstrate the capabilities of the Polarity-JaM pipeline, by quantifying the intensity ratio of Krüppel-like factor 4 (KLF4) in the nucleus with respect to the cytosol. KLF4 is a transcription factor that is known to be upregulated via exposure to laminar shear stress (49, 50). We calculated the intensity of KLF4 in the nucleus and cytosol for static, after 4 h and 16 h of 6 dyne/cm^2^ flow. We found a significant increase after 4 h of flow exposure in nuclei localisation compared to control and a slow decrease at 16h compared to 4h Fig. 5 B,C. For statistical comparison, we have used the DABEST method (51), see Fig. 5 C.

**Fig. 5.**
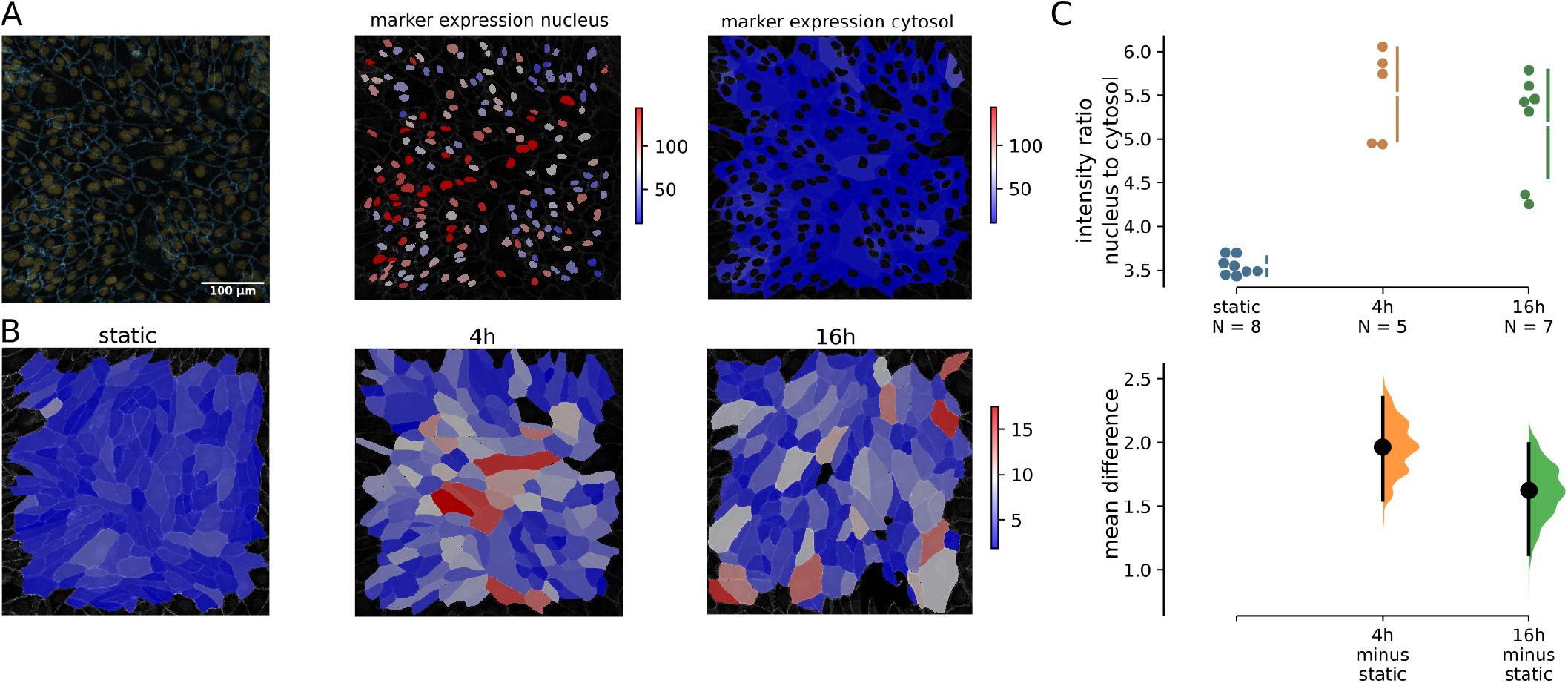
Localised single-cell fluorescence intensity quantification. **A:** Quantification of the intensity of KLF4 antibody staining. Overlay KLF4 and junction channel [left], KLF4 intensity in the nuclei [middle] and in the cytosol without nucleus [right]. **B:** Intensity ratio of the KLF4 reporter in the nucleus and in the cytosol was colour coded for each single cell. **C:** Statistical comparison of the change using the DABEST method shows a significant difference between static conditions and endothelial cells exposed to flow after 4 and 16 h. Each dot indicates a different image with N=50 - 100 cells each. Black bars indicate 95 % confidence intervals of the mean difference.

### Junction morphology

Cell–cell junctions underpin any architecture and organisation of tissue. They vary in different tissues, organs, and cell types and need to be dynamically remodelled in development, homeostasis, and diseases. For example, endothelial cell-cell junctions must provide stability and prevent leakage while also allowing dynamic cellular rearrangements during sprouting, anastomosis, and lumen formation (52). The organisation and topology of junctions and inversely the organisation and topology of endothelial cells contain a wealth of biological information. By analysing adjacency patterns in endothelial cells, organisational patterns that are associated with tissue phenotypes can be uncovered. There are vast differences in endothelial arrangement between different tissues and organs (53).

We are using the cell–cell contact features from an already published tool JunctionMapper to decipher cell-cell junction-related phenotypes (20). Note, that this tool is not adapted to studying EC in organs or in 3D tubular structures, which will be in the scope of future studies. The normalised junction features suggested in JunctionMapper allow one to quantify images of different resolution, cell type, and modalities. In our tool, the analysis is automatically performed in a region that is defined by the cell outlines, which we obtain from the instance segmentation using the proposed deep learning frameworks. This outline is dilated by a user-defined thickness. There are no more parameters necessary to define. The result can be seen in Fig. S1 C. The resulting area of the dilated outline is the interface area, which is computed for each single cell. We then derive the characteristic from a junction label, for example, VE-cadherin staining in the case of ECs, see Fig 6 A. The fragmented junction area results from Otsu thresholding in that region, see Fig. 6 B. Using these readouts, we can compute three features: (1) interface occupancy by computing the fragmented junction area over the interface area, (2) the intensity per interface area by computing the average intensity in the interface area, and (3) cluster density by the average intensity in the fragmented junction area. We find a unique signature of the three junction features in ECs after flow stimulation, see Fig 6 C, demonstrating the effectiveness of our method. In static condition, the cell-cell junctions are very heterogeneous, with some cells having thick junctions and high VE-cadherin intensity, while others have low signal intensity and low occupancy. Intensity and occupancy become more homogeneous within each field after exposure to flow. At 6 dyne/cm^2^ the total intensity per interface area increases as well as the interface occupancy. At 20 dyne/cm^2^, however, the junctions become thinner, resulting in lower interface occupancy, while the intensity per interface area remains almost the same compared to static. At the same time, the intensity within the junction increases, resulting in higher values of cluster density. For the entire junction analysis workflow, only one additional parameter needs to be specified, namely the width of the automatically generated outlines that serve as regions of interest for cell-cell contacts. In summary, Polarity-JaM offers the possibility to fully automate the essential parts of the JunctionMapper workflow by setting a single additional parameter.

**Fig. 6.**
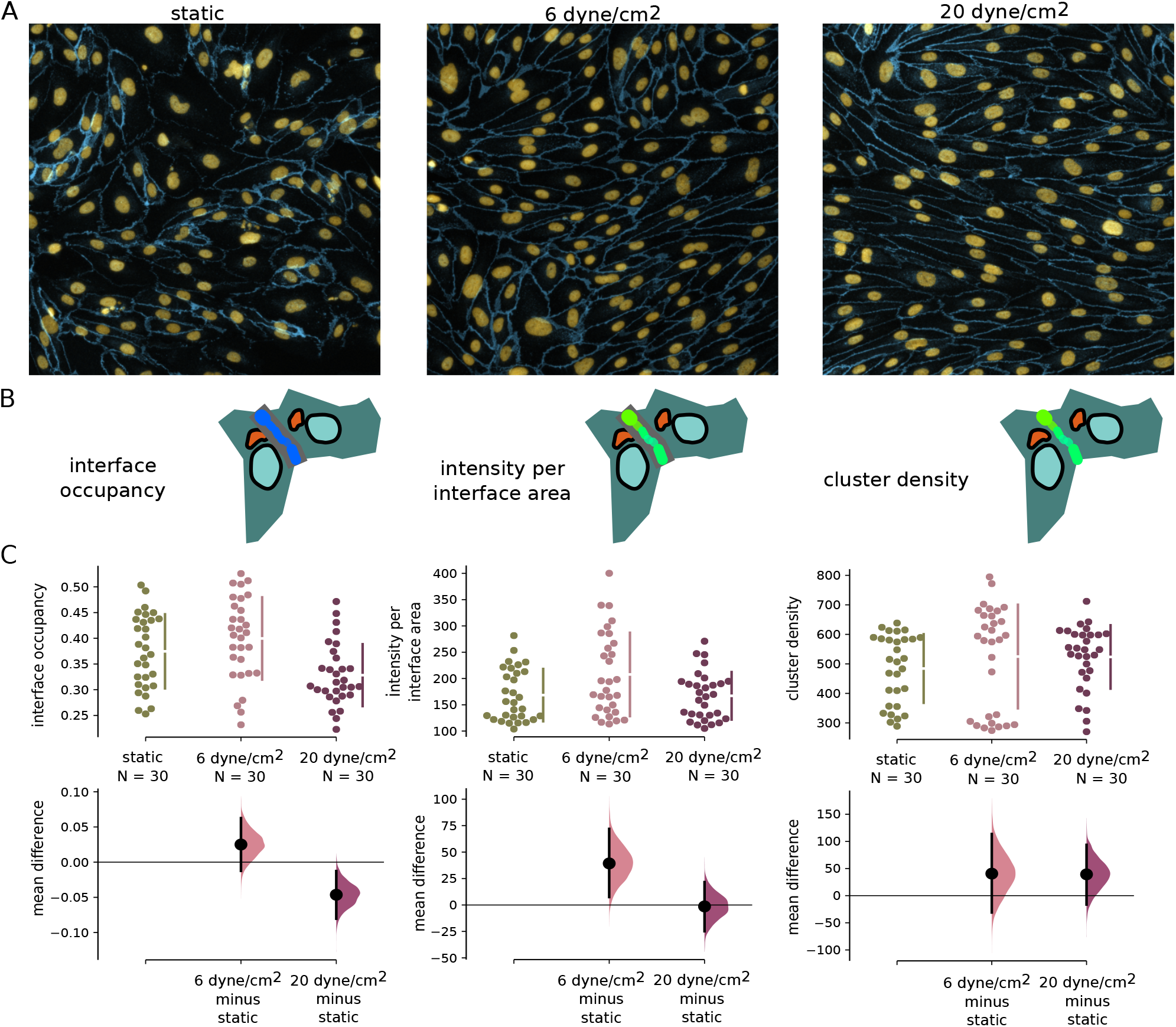
Junction morphology quantification. **A:** Examples of endothelial cells in static condition, exposed to 6 dyne/cm^2^ and 20 dyne/cm^2^. VE-cadherin staining is shown in blue. **B:** Pictograms of quantities extracted from the images. **C:** Statistical comparison of three normalized morphological junction features interface occupancy [left], intensity per interface area [middle] and cluster density [right] demonstrate a significant change in junction dynamics after exposure to flow. Black bars indicate 95 % confidence intervals of the mean difference.

### Reproducability, Replicability and Interoperability

Iterative acquisition of images and various experimental settings sometimes require complex folder structures and naming schemes to organise data, leaving the researcher with the problem of data structure and reproducibility of their analysis. To help with both tasks, the Polarity-JaM pipeline has three execution scenarios: a) single image, b) image stack, and c) complex folder structure. Furthermore, a comprehensive logging output is provided, as well as a standardised input structure in yml format. The generated outputs follow a naming scheme. The extracted collective and single cell features are stored in a csv file. The results of the statistical analysis from the app can be downloaded in various formats, including pdf and svg. For different categories or conditions, the Polarity-JaM app uses several qualitative colour schemes that are colour-blind friendly (54), the infrastructure follows a similar principle as PlotTwist, which was designed for time series analysis. Metadata and log information are also saved in a human-readable format on disk. Polarity-JaM uses well-established non-proprietary formats (such as csv, yml, tiff, svg) to aid interoperability, following a recommendation in (55). All statistical analysis for circular features shown in this study and more can be performed in the App. Our tool can be combined with other tools such as Griottes (21), polarity features can be mapped on spatial network graphs and their relation can be explored using the same segmentation.

Exploitative image analysis requires interactivity to quality check each analysis step. Hence, Polarity-JaM is designed with a simple Python API that is optimised for usage within a Jupyter Notebook(56). We provide several examples in our documentation on how to perform such an analysis. An overview of the entire Polarity-JaM software suite is depicted in Fig. 7. We additionally equip Polarity-JaM with a Napari (57) plugin with a graphical user interface to enable direct feedback on segmentation and features. Finally, Polarity-JaM is available via PyPI (Python Package Index). Taken together, we are committed to the principles of FAIR research (58).

**Fig. 7.**
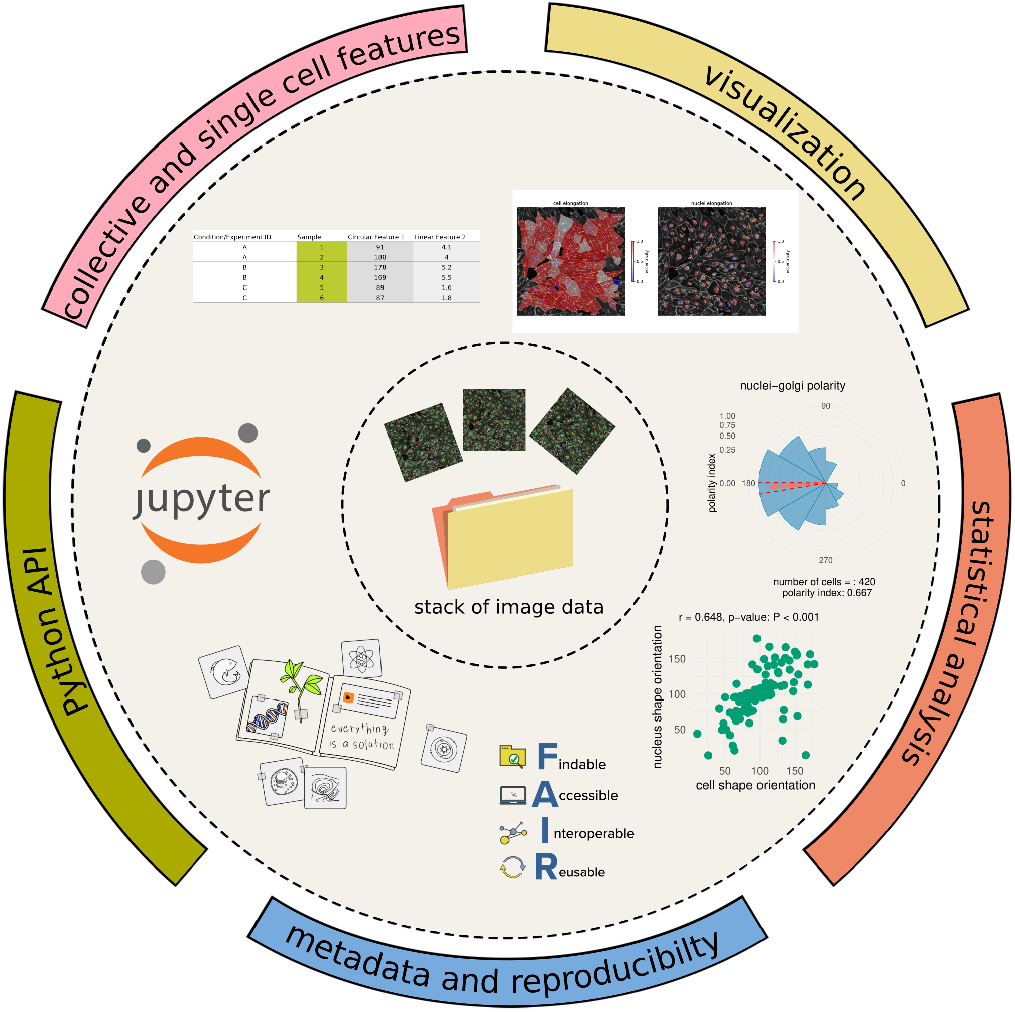
The workflow enables processing large image data stacks for automated feature extraction. The Python API can be used to facilitate quality control with the help of a Jupyter Notebook. The results can be analysed statistically using a graphical user interface.

## Discussion

Our image data processing and analysis workflow can be used to simultaneously compute features of cell polarity, including organelle localisation, cell shape, and signalling gradients, allowing single-cell and collective high-content endothelial phenotyping. Circular statistics can be performed interactively via a web application. We also provide an informative graphical design for directional and axial data. We recommend the use of the signed polarity index when a polarisation direction is expected on the basis of an external polarisation cue, as is the case in most endothelial flow assays.

With the focus of the Polarity-JaM toolbox on a diverse feature set and replicability and interactivity, we provide means for answering various biologically motivated experimental questions and for the extraction of aspects that are otherwise easy to overlook. For instance, collective orientation and cell size have been inversely correlated with endothelial cell senescence. Older endothelial cells tend to be larger and share a less pronounced direction of orientation (37). The orientation as a collective phenotype can be correlated with processes such as flow and senescence. Multivariate analysis, for example, including size in the image analysis, could perhaps differentiate between endothelial phenotypes by flow or those by cellular senescence.

This investigation has been exemplified for endothelial cells, but can also be applied to other cell types such as the planar polarity of cardiomyocytes (59), epithelial cell tissues (19) and potentially other cell types. The image modality is not restricted to fluorescence microscopy but can be also applied to phase contrast or other here the only restriction is that the image can be decomposed into masks of single cells. The current version of Polarity-JaM integrates different segmentation models, including Cellpose (8), microSAM (7) a fine-tuned model based on SAM (6), and DeepCell (5). There is a vivid community around all these segmentation algorithms (7, 8); therefore, we provide an interface to these models, which can also be adapted by the user. This will help maintain this software and ensure long-term use. Future development needs to address better segmentation of subcellular structures, including cell-cell junctions, cytoskeleton, and focal adhesion sites, using deep learning methods.

The quantification of junctional morphology is based on the features suggested in (20) including junction occupancy, cluster density, and intensity per interface area. While these features provide good indicators for junctional changes and adaptations, they may not be exhaustive referring to the manual morphological classification of adherens junctions, which is frequently done in five common categories: straight junctions, thick junctions, thick to reticular junctions, reticular junctions, and fingers (60). To automate the translation into this classification, further work is needed on junction segmentation, as well as an advanced classifier using manual training data, which was not ready at the time of this publication but will be addressed in the future.

Future challenges involve tissue and organoid image data in 3D space, which introduces more challenges in algorithmic development including robust segmentation (mainly due to the lack of training data), anisotropy in image acquisition, and the size of the image data. Also, efficient extraction of cell and nuclei features, which are by default not included in common packages, need to be developed. Multiplex imaging will stimulate further developments, as this image data modality dramatically increases the content of the information and therefore challenges meaningful feature extractions and comprehensive statistical spatial and circular approaches to compute cross correlations (61).

The focus of this pipeline is on static images. However, the pipeline could also be applied to a series of images, and feature extraction would be performed for each frame. The extracted data can be stitched together by label identification (62). Computational models can be informed by a wealth of quantitative data through our approach, including vertex models (40), but also the cellular Potts model(35, 63) or agent-based models (64, 65). The spatial context of can be further explored using tools such as Griottes (21) and the circular version of Moran”s I (66) to extract collective phenotypes. Mechanistically, this will also help to predict different states of tissues and dynamics from static biomedical images (41, 67, 68), which is an interesting avenue for future research with a wide range of applications.

## Materials and Methods

### Experimental setup

#### Cell culture and fluid shear stress assays

Commercially available human umbilical venous endothelial cells (HUVECs, pooled donors, PromoCell) were cultured and used at passages 2-4 at 37°C and 5% CO_2_ humid incubator, in endothelial growth medium containing growth factors (EGM2, Lonza) for optimal cell growth. For passaging and fluid shear stress (FSS) assays, cells were washed once in sterile PBS, followed by a 5 minute incubation in Trypsin at 37 °C and 5% CO_2_, then neutralized with FBS and EGM2. Cells were centrifuged for 5 minutes at 480 g and counted. For FSS assays, cells were seeded in 0.4 ibiTreat Luer flow slides (Ibidi) coated with 0.2 percent gelatin at a cell concentration of 2 million cells per ml. 100 µl of cell suspension were added to each slide and incubated overnight at 37°C and 5% CO_2_. The following day, the slides were connected to red perfusion sets assembled onto perfusion units (Ibidi) and connected to a pump (Ibidi). Laminar shear stress was applied at 6 dyne/cm^2^ or 20 dyne/cm^2^ for 4, 16, or 24 h inside a 37 °C and 5 % CO_2_ humidity incubator. Static controls were kept in the same incubator for the duration of the experiment.

#### Immunofluorescence

At the end of a FSS assay, slides were disconnected from perfusion units and immediately fixated in 4 % PFA for 10 minutes, then washed 3 times in PBS. Slides were then blocked and permeabilized in CBB (Claudio”s Blocking Buffer, see supporting information) for 30 minutes, followed by 1 h of incubation with primary antibodies at room temperature. Cells were washed 3 times with PBS, then incubated 1 h with secondary antibodies at room temperature, followed by another triple washing step, 5 minute incubation with DAPI, and finally mounted with a Mowiol + Dabco mixture in a 9:1 ratio. The primary and secondary antibodies used can be found in Table S1.

#### Confocal imaging

FSS immunostained slides were imaged on a confocal microscope (Carl Zeiss, LSM 980) using a Plan-Apochromat 20x/0.8 NA Ph2 air objective. For each sample, 10 random positions throughout the flow slide were be selected and a Z-stack covering the entire depth of the monolayer were acquired. Slides were imaged with a two-channel setup, with channel one using the 488 and 633 lasers and channel two using the 405 and 561 lasers. Pinhole size was set to 1AU for both channels.

### Image Analysis

#### Segmentation

To isolate individual cells in a microscopic image, a process also known as instance segmentation, we used Cellpose, a deep neural network algorithm. Accurate instance segmentation can be created with pre-trained models it is provided with. These models can generalise well across both cell type and image modalities. For our analysis we used the model “cyto” for cell and “nuclei” for nuclei instance segmentation. For Golgi segmentation, Otsu-thresholding was performed. Subsequently, the segmentation mask was used to get the corresponding Golgi instance label. The performance of instance segmentation algorithms can vary for different modalities. Downstream analysis of features describing individual cells and their relationship with each other strongly depend on the quality of these segmentations. At this point Polarity-JaM offers three segmentation algorithms that the user can choose from: Cellpose, DeepCell, and microSAM. Additional segmentation algorithms are realised with the help of Album (69), a decentralised distribution platform where solutions (in this case implementations of segmentation algorithms) are distributed with their execution environment and can be used without additional overhead for the user. All of these show state of the art performance and are included in Polarity-JaM to enable segmentation for a broad range of modalities and cell types.

#### Single cell and organelle features

Common features are available within the *regionprops* scikit-image package (70). We extend the available measurements by various features. For a complete list of all features, see the Appendix 3. Most features require central image moments (71) that can be calculated from the raw moments

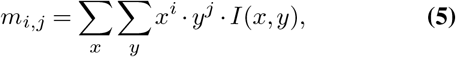

with *i, j* = 0, 1… are exponents, x,y the pixel coordinates, where *I*(*x, y*) refers to the image intensity at position x,y. Generally, the centre of mass of a grey scale image (e.g. a channel) is now given by

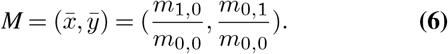

The central moments are then

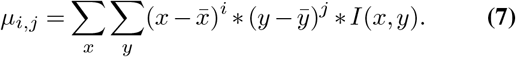

#### Shape orientation

With the central moments, we compute the orientation

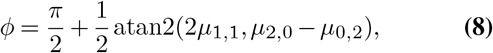

which describes the angle of the major with the x-axis in the interval [0, *π*] in radians or 0 to 180 degrees. Various features can be defined with the orientation, such as the cell shape orientation or the nucleus orientation in case the nucleus channel is provided.

#### Nucleus and organelle displacement

The displacement orientation from the nucleus to the centre of mass of the cell can be defined as

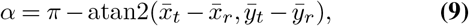

where index *r* indicates moments that are calculated on a reference channel, and *t* moments that are calculated on the target channel. In the case of nuclei-Golgi polarity, the target is the Golgi channel and the reference the nuclei channel. The orientation takes values in [0, 2*π*] in radians, which corresponds to 0 to 360 degrees, in this study. Analogously, the orientation from i) nucleus to organelle, ii) centre of mass of a marker channel to cell centroid, and iii) centre of mass of a marker channel to nucleus centroid can be defined.

#### Signaling gradient quantification

We define the

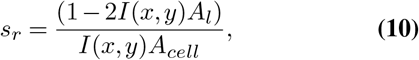

for a cell, where *A*_*l*_ is the area of the left cell half perpendicular along a given a cue direction *θ* and *A*_*cell*_ the area of the cell. Mathematically, the area *A*_*l*_ is described as

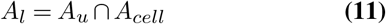

with

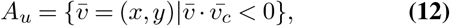

And 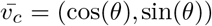, where the polar direction or expected cue direction *θ* is given in radians.

#### Circular statistics

All statistical methods were implemented in R and R-shiny. We have applied the statistical tests from the CircStats package (31) for directional and axial data. For measures on a linear scale, we applied the DABEST (“data analysis with bootstrap-coupled estimation”) method from (51).

### Availability, code, and issues

The pipeline was developed in Python and is available through PyPi(72). The code for the R-shiny(73) application was written in R (https://www.r-project.org) and Rstudio (https://www.rstudio.com). The Polarity-JaM app can be used online via (www.polarityjam.com) without the installation procedure. Both the Polarity-JaM pipeline and the app can also be used offline. Additionally, both can be installed and used separately through album (69), a framework for scientific data processing with software solutions of heterogeneous tools. The link can be found https://album-app.gitlab.io/catalogs/helmholtzimaging/de.mdc-berlin/polarityjam. The code for pipeline, app, and Napari plugin is published under the MIT licence and is available through GitHub (https://github.com/polarityjam). Additional documentation and information can be found at readthedocs (https://polarityjam.readthedocs.io). Issues and requests are tracked on GitHub issues. Collaboration and extension are possible and welcome. Instructions and best practices can be found in the documentation.

## ACKNOWLEDGEMENTS

The authors thank the IT department of the MDC, especially Frank Büttner for his help and support in deploying the R-shiny web application on the MDC server. In questions of data protection and the legal framework for the online application, we were supported by Ulrike Ohnesorge and thank her for her commitment. For this bioRxiv, we used the https://henriqueslab.github.io/ resources/bioRxivTemplate/ template by Ricardo Henriques from the Francis Crick Institute. The authors also thank Christoph Karg for valuable feedback.

## AUTHOR CONTRIBUTIONS

**Wolfgang Giese**: Conceptualisation, Data Curation, Formal Analysis, Investigation, Methodology, Project Administration, Software, Supervision, Validation, Visualisation, Writing – Original Draft Preparation, Writing – Review & Editing; **Jan Philipp Albrecht**: Conceptualisation, Data Curation, Formal Analysis, Investigation, Methodology, Software, Validation, Visualisation, Writing – Original Draft Preparation, Writing – Review & Editing; **Olya Oppenheim**: Data Curation, Formal Analysis, Investigation, Validation, Writing – Original Draft Preparation, Writing – Review & Editing; **Emir Akmeriç**: Investigation, Methodology, Writing – Review & Editing; **Julia Kraxner**: Investigation, Visualisation, Writing – Review & Editing; ; **Deborah Schmidt**: Funding Acquisition, Resources, Writing – Review & Editing; **Kyle Harrington**: Conceptualisation, Funding Acquisition, Methodology, Supervision, Visualisation, Writing – Review & Editing; **Holger Gerhardt**: Conceptualisation, Funding Acquisition, Resources, Supervision, Writing – Review & Editing.

## Supplementary Note 1: Figures

**Supplementary Figure S1.**
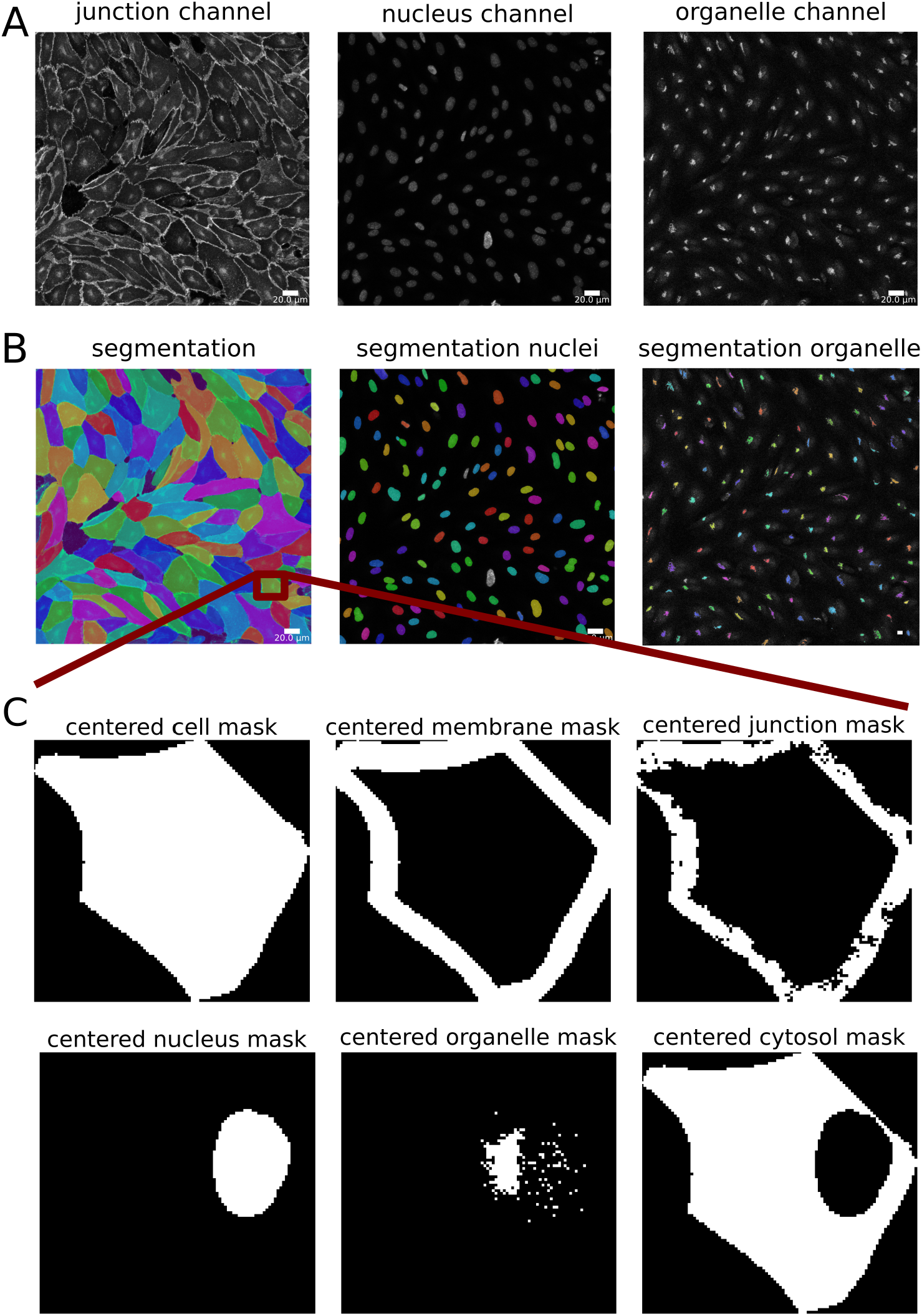
Segmentation visualizations. **A:** Channel information **B:** Cellpose cell and nuclei segmentation, otsu threshold on organelle channel (here: Golgi) **C:** View on a single cell, together with the membrane, junction, nuclei, organelle (here: golgi) and cytosol mask.

**Supplementary Figure S2.**
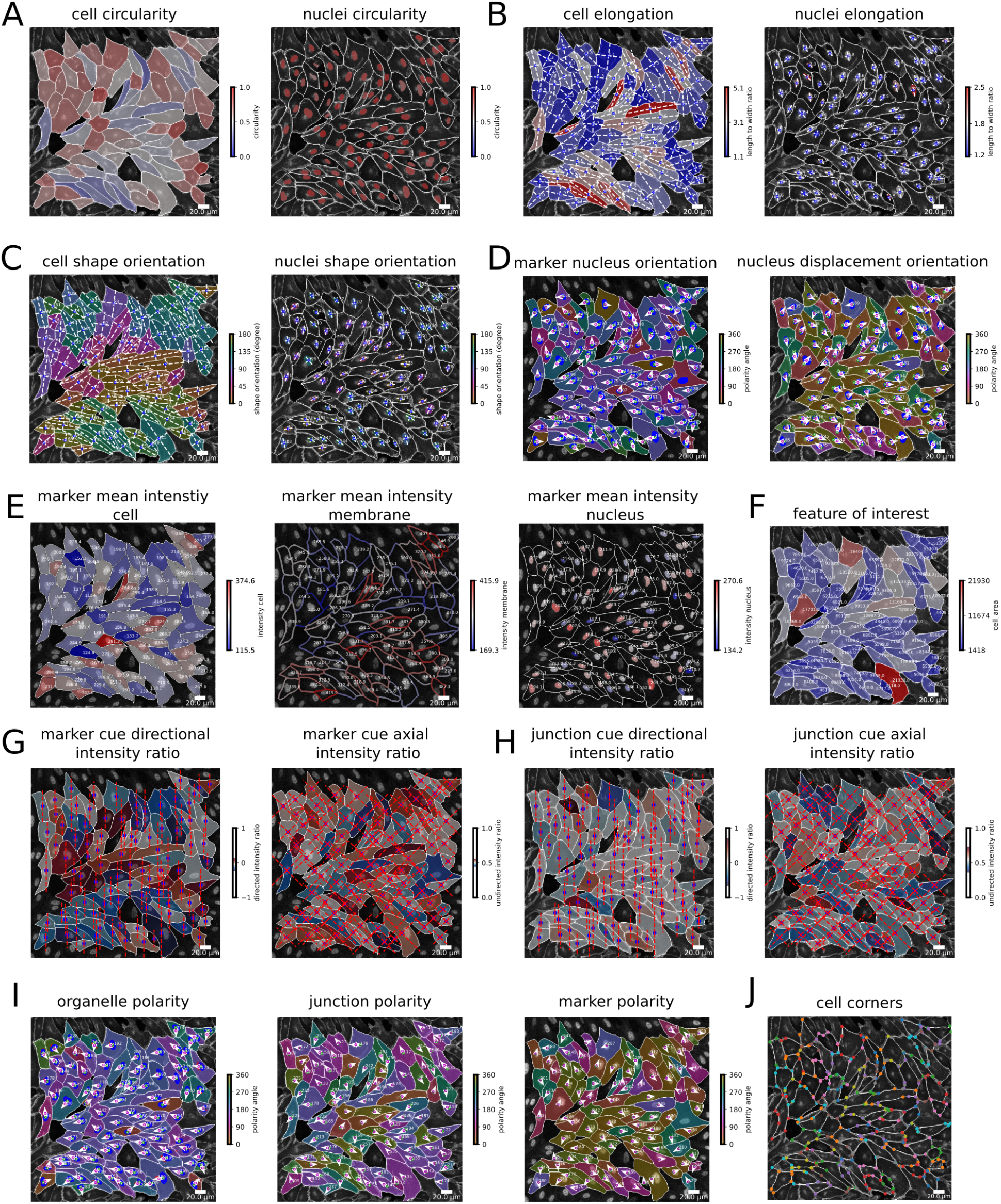
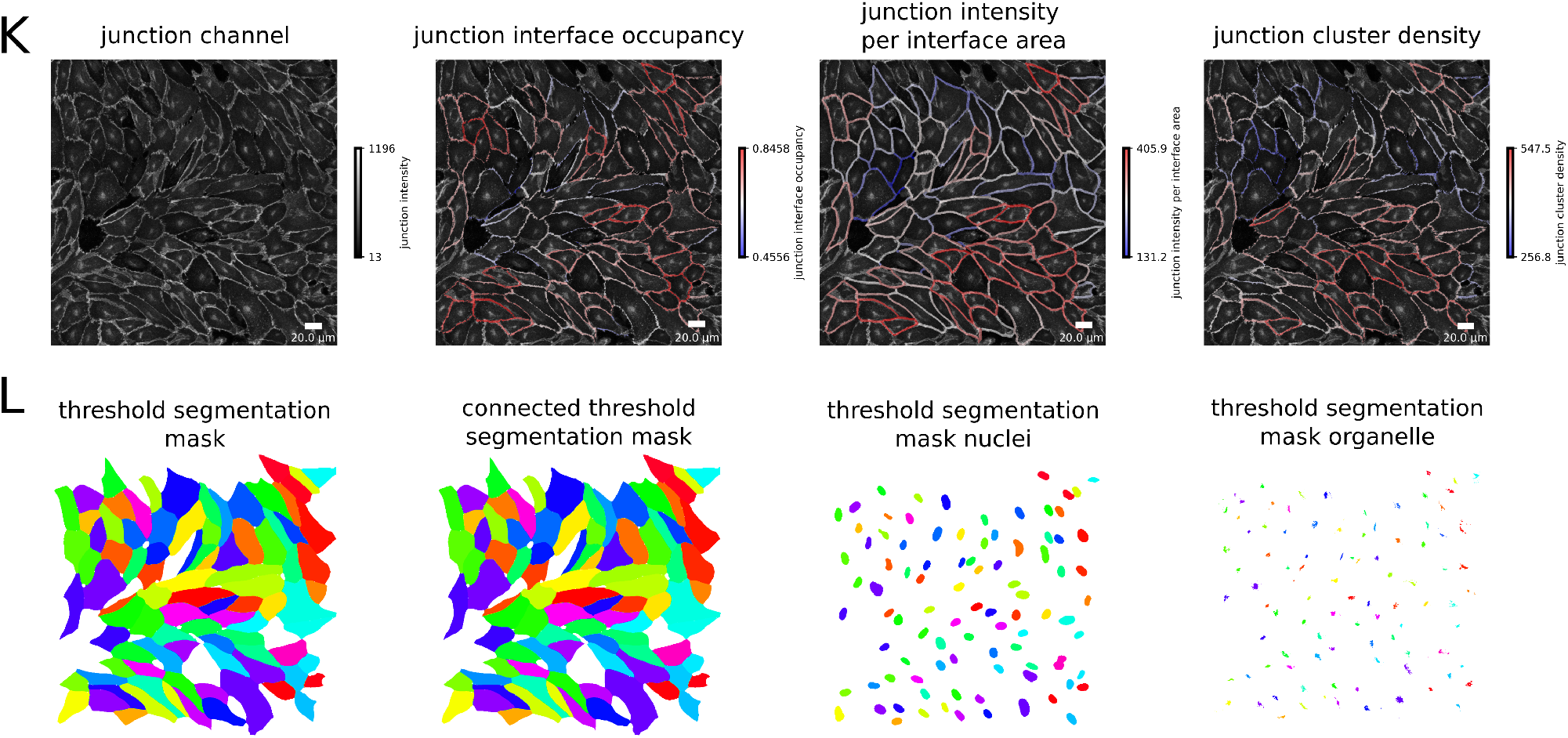
Feature visualizations. **A:** Circularity, **B:** Elongation, **C:** Shape orientation, **D:** Marker nucleus and nucleus displacement, **E:** Intensity information, **F:** Feature of interest (here: area) information, **G:** Marker ratio method, **H:** Junction ratio method, **I:** Polarity information for organelle(here: golgi), junction and marker channel, **J:** Cell corners based on the Douglas-Peucker-Algorithm. Feature visualisations continued. **K:** Junction features, including channel, interface occupancy, intensity per interface area, and cluster density, **L:** Segmentation masks after applying a threshold for cell, nuclei, and organelle size.

**Supplementary Figure S3.**
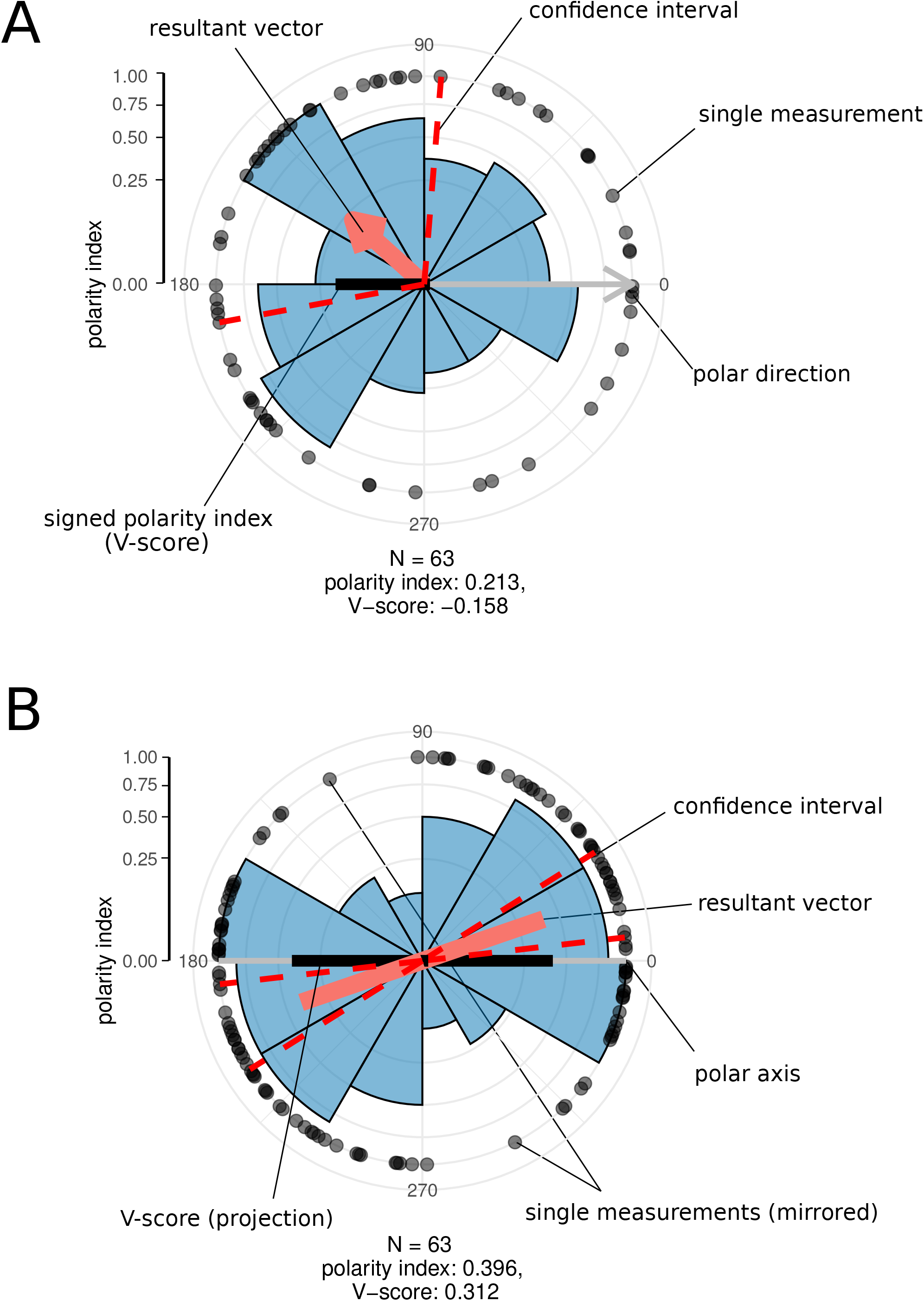
Circular statistics data graphics with suggested statistical measures for **A** directional data and **B** axial data.

**Supplementary Figure S4.**
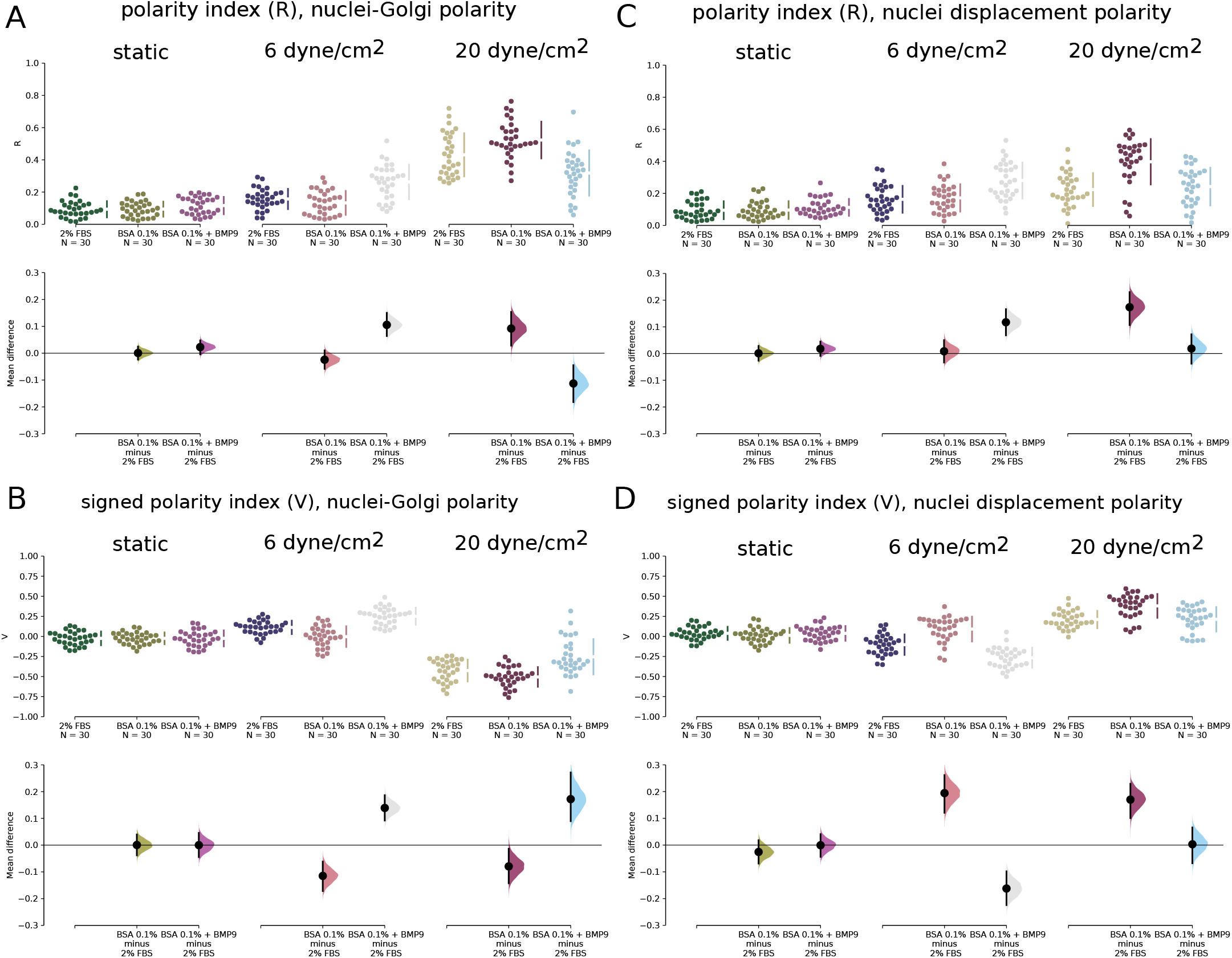
Circular statistics data graphics that compare polarity index (R) and signed polarity index (V) for all conditions shown in Fig. 2. Black bars indicate 95 % confidence intervals of the mean difference.

**Supplementary Figure S5.**
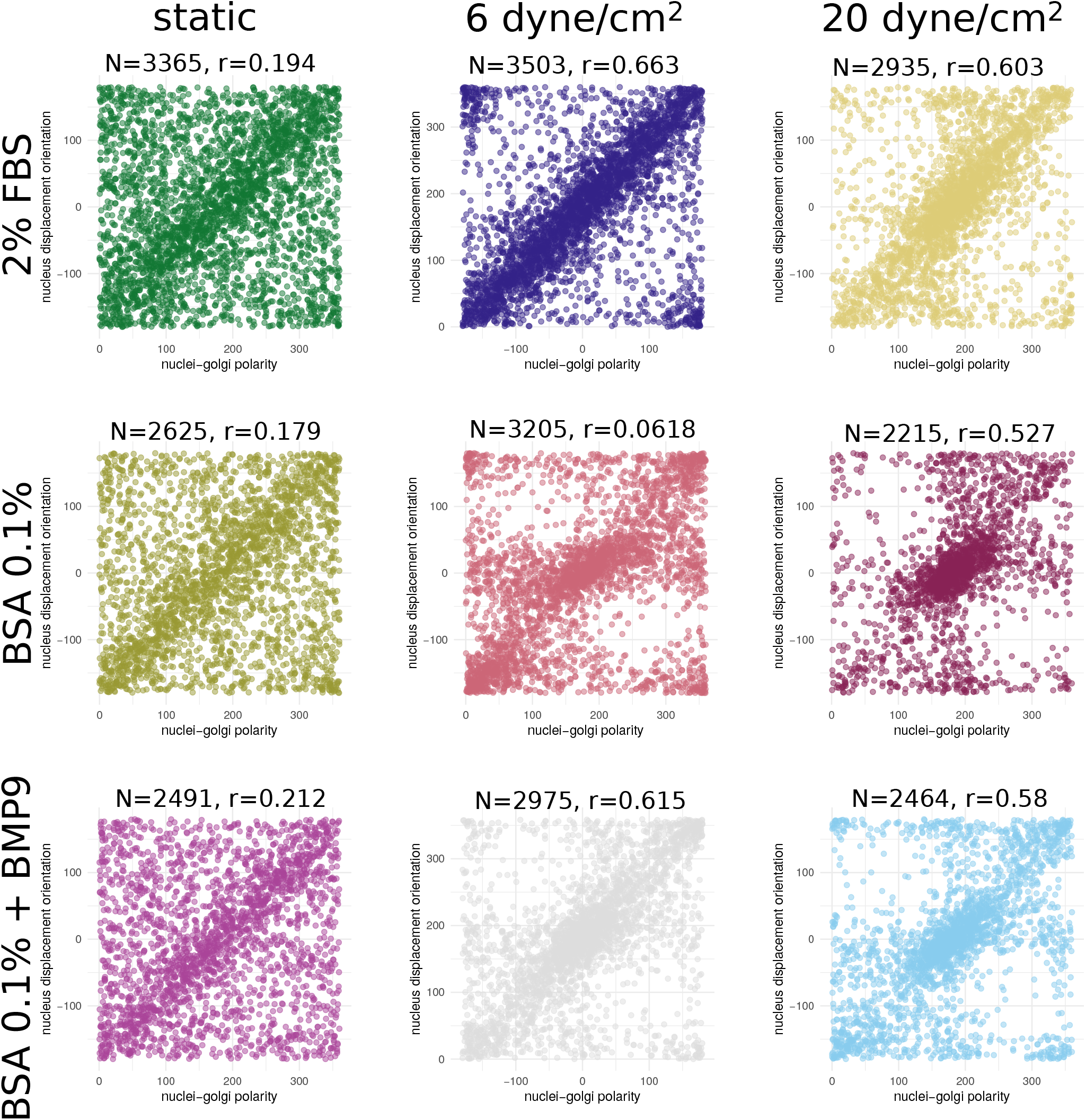
Circular correlations of nuclei-Golgi polarity and nuclei displacement for different flow and media conditions corresponding to Fig. 2.

**Supplementary Figure S6.**
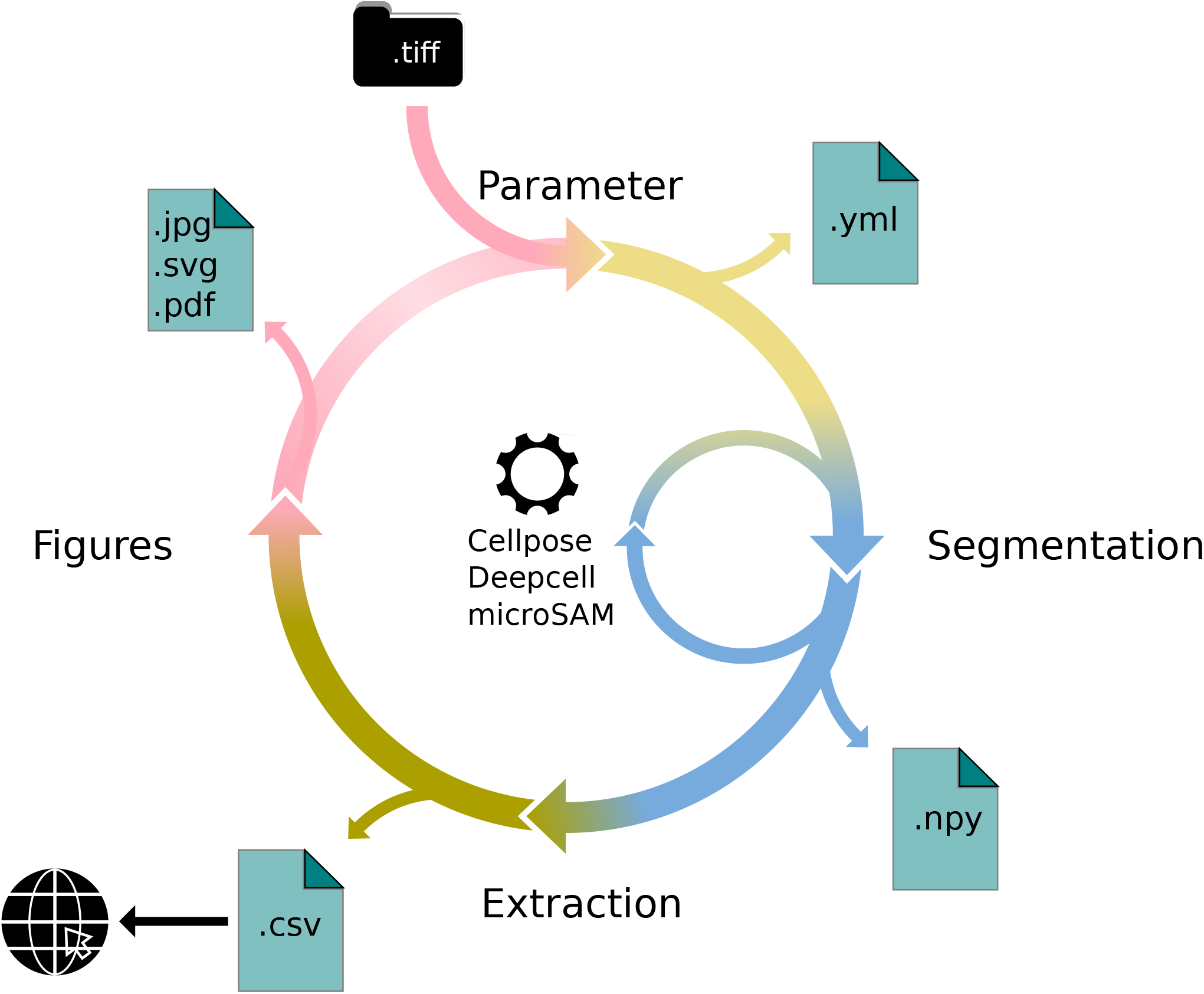
Visual depiction of the python API. After loading the image, the user would first define parameters describing the details of the analysis. Next step would be the instance segmentation. Currently three algorithms, Cellpose, DeepCell, and microSam are supported. After the feature extraction process the visualization functionality of Polarity-JaM could be used to asses data quality.

## Supplementary Note 2: Data input and structure

Often, analysts are challenged not only with the problem of actually performing the analysis, but also how and where to store the data. Iterative acquisition of images as well as various experimental settings sometimes require complex folder structures and naming scheme to organise data. Frequently, researchers face the problem of data being distributed over several physical devices, leaving them with the problem of how to execute a certain tool on a dedicated subset of images. Often a large amount of time is spent organizing data before the analysis can finally be conducted. Moreover, performing analysis steps on several experimental conditions often requires repeating the entire analysis pipeline several times to get the desired output. Another problem researchers often face is keeping track of the configuration that was used for a particular execution of software at a certain time. Sometimes this even requires some form of manual work by writing down the configuration. Polarity-JaM avoids these unnecessary chores and manual processes for the user and minimizes the time spent for reorganizing data by providing a) three run cases to cover most usage scenarios, b) a parameters yml file containing all options used for the analysis, and c) a log file fully capturing the output of the analysis.

### Parameters

The parameters file is the most important argument that needs to be passed to an execution call as it specifies what analysis steps are performed and hence what output will be created. The file additionally is copied to the output folder and hence allows the user to fully reproduce and comprehend the analysis at a later time-point. An overview of all possible keywords is given in Table S2. This includes configuration parameters describing the image (image parameter), analysis (runtime parameter), and visualisation (plot parameter). Please note that the parameters necessary for the segmentation algorithm are listed separately in Table S3.

### Run Options

To tackle the problem of various examples of use, Polarity-JaM offers three run options. The mode *run* can be used for a single image and only requires an input image, and an output path. *run-stack* should be chosen if a set of images in a folder needs to be processed. Instead of a single image, a folder can be specified. Lastly, the *run-key* option needs a csv file describing the storage structures and experimental conditions and must be passed as an argument. The structure of the csv is deliberately kept simple and is shown in table S4. Paths are relative to an input path that can be passed separately as an argument to the execution call. Hence, the data can be copied to a different physical device and the analysis can be repeated without altering the csv file as long as the underlying data structure remains untouched.

## Supplementary Note 3: Features

The feature extraction pipeline is the process of extracting all relevant features from an input image. This is a complex process that can be separated in three major parts. First, the image is segmented, segregating each cell such that in the second step its features can be extracted. In the third and last step, a graph structure is build and a neighborhood analysis performed. To the time writing the manuscript, the user can choose between three segmentation algorithms: Cellpose(8), microSAM(7) a fine-tuned model based on sam(6), and DeepCell(5).

The pipeline produces various outputs, depending on the parameter configuration and input provided. In general, generated output can be differentiated in the two categories a) features, and b) visualization. Features are gathered in a csv file containing the individual cells as rows and their corresponding feature values as columns. Visualizations for the extracte features can be optionally created and written to disk. These plots should be used for quality control before continuing downstream analysis of the extracted features. An example of all available visualizations is shown in figure S2.

### Categories

The features can be structured in several categories which will be explained shortly. Depending on the configuration, the categories will be included or excluded in the extraction process. The categories are shown in table S5. Each cell has various pieces of general information such as area, perimeter and elongation, which are summarised in the *single cell features* that are shown in table S6. The category *neighborhood features* contains all properties that can be addressed to the surrounding of a cell. An overview is found in table S7. Given a feature of interest (e.g. cell size) the Morans I correlation analysis can be performed. *group_features* are depicted in table S8. Please note that their values address an entire image and not a single cell and should be interpreted accordingly. Optionally, *nucleus features, organelle features, marker features*, and *junction features* can be gathered and are explained in tables S9, S10, S11 and S12 respectively. Nucleus features mainly comprise morphological features, such as their size, position, orientation. Organelle features contain positional, but also distance information to the nucleus, whereas marker features include besides positional, also mean expression values of several cell regions. (e.g. membrane, nucleus and cytosol). Junction features mainly comprise ratio values between different areas of the junction region of the cell membrane (20), but also mean intensity information and polarity.

## Supplementary Note 4: Visualizations

Beside features, several visualizations are created during the analysis. They mainly serve as quality control. To the point of manuscript writing Polarity-JaM provides 18 different visualizations:

- image channel intensity information
- segmentation masks
- threshold segmentation masks
- single cell closeup
- cell corners
- feature of interest
- cell and nuclei elongation
- cell and nuclei circularity
- cell and nuclei shape orientation
- nucleus displacement orientation
- organelle polarity
- nucleus marker orientation
- marker polarity
- marker expression
- marker cue intensity ratio
- junction features
- junction polarity
- junction cue intensity ratio

Quality control of the segmentation output is a crucial step for downstream analysis. For this purpose, the feature extraction pipeline creates for every input image a plot showing the given channel configuration used for segmentation together with the segmentation outcome. (Figure S1 A,B)). Optionally, a closeup view for each cell can be plotted as shown in figure S1 C. Feature visualizations are shown in figure S2 And highly depend on the quality of the segmentation. We highly recommend to perform a quality control of the segmentation result before continuing downstream analysis for example with the Polarity-JaM web app.

## Supplementary Note 5: API Usage

Polarity-JaM offers the possibility to completely run an anaylsis within a Jupyter Notebook (74). Several example notebooks can be found in the GitHub repository https://github.com/polarityjam/polarityjam/tree/main/docs/notebooks focusing on various aspects of an analysis. This includes the basic usage of Polarity-JaM to perform a feature extraction to using Polarity-JaM as a python library for enhanced image analysis. We now shortly describe a basic analysis in chronological order. First, the user loads the image. Second, the user sets up the parameters for i) the image(s), ii) the runtime (analysis procedure), including the choice of the segmentation algorithm, and optionally iii) the visualizations. Additionally, parameters will be saved on disk in yml format to support replicability. Third, the previously set segmentation algorithm is loaded, initially with its default parameters. At this point the user has the option to alter these before performing the segmentation. Any segmentation algorithm that is currently supported in Polarity-JaM requires two steps for the user to perform: i) preparing the image for segmentation, and ii) using the prepared image to perform the segmentation. The division in two steps is specifically designed to facilitate visual quality inspection by using the visualization functionality of Polarity-JaM. Additional segmentation algorithms (other than Cellpose) are implemented with the help of Album (69), a decentralized distribution platform where solutions (in this case implementations of segmentation algorithms) are distributed with their execution environment. This allows to easily switch to a different segmentation algorithm when performance is poor. The overhead of installing the algorithm in its correct environment is taken from the user. Regardless of the used algorithm the analysis steps follow the same semantic (e.g. the same function call) such that usage of an unknown algorithm is facilitated. Output of a segmentation procedure is always an instance segmentation mask in numpy (75) format. We provide detailed information in a Jupyter Notebook that can be found under the following link: https://github.com/polarityjam/polarityjam/blob/main/docs/notebooks/polarityjam-notebook_seg.ipynb To improve downstream analysis, instance masks for nuclei and organelle should additionally be calculated whenever the corresponding information is present in the image. To facilitate the process for the user, every segmentation algorithm supported in Polarity-JaM has a mode that can be specified in the segmentation step. Currently, there are three modes available: “nucleus”, “cell”, and “organelle”. Please note that not every algorithm supports all modes. The user is however free to entirely skip any segmentation step and provide instance segmentation masks elsewhere produced. As a fourth step, the user performs the actual feature extraction by first setting up an initially empty collection that can be then passed to the routine performing the extraction, together with the instance mask(s), the original image, and the parameter describing the image. Gathering features in a collection allows the user to potentially collect features of various images for example by looping over images in a given folder. Last step should always be to use the visualization functionality of Polarity-JaM to assess quality and get a first impression of selected features before moving on to the downstream analysis of the features in the Polarity-JaM Web-App. The entire API is depicted in Figure S6 And shows the workflow the user performs when working in a Jupyter Notebook.

## Supplementary Note 6: Circular statistics

A broad range of scientific studies involve taking measurements on a circular, rather than a linear scale (often variables related to orientations or circadian times). However, their analysis is not straightforward and requires special statistical tools. All features in our study were classified into periodic and linear features. Among periodic features, we further distinguish directional features with values in a full circular scale, meaning that the data repeat after 360 degrees (or 2 *π* in radians) and axial features that repeat after 180 degrees (or *π* in radians). Axial data refer to an axis, in our case the long axis of the cell or nucleus shape, rather than to a direction.

Circular data presents some unique challenges for statistical analysis because traditional statistical methods may not be appropriate for this type of data. For example, computing the average or mean of circular data by summing the values and dividing by the number of observations can produce incorrect results. Therefore, specialised statistical methods have been developed to analyse circular data, such as circular statistics and directional statistics. These methods take into account the periodic nature of the data and can provide a more accurate result. All the different features such as nuclei-Golgi polarity, cell shape orientation, cell elongation, or junction properties can be correlated amongst each other compared and correlated to linear read-outs such as cell size or cell elongation on a single cell level.

Several of the extracted polarity features are periodic and require different means of statistical comparison. The polarity index is defined as follows

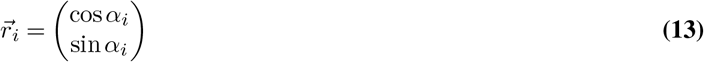

and the average resultant vector

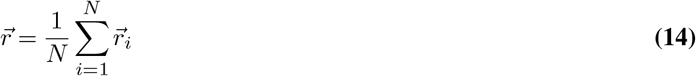

Although the polarity index describes the concentration of the distribution with respect to the mean direction, we may also be interested in a measure that quantifies the degree of orientation towards a given direction *µ*_0_, which is referred to as the polar direction (31). This is described by the V-score, which is computed from

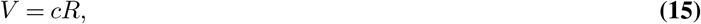

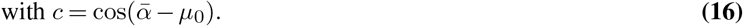

Therefore, *V* is equal to the polarity index, *V* = *R* if aligned with the given direction and takes the negative value if aligned with the given direction *V* = *−R*.

These statistical measures can also be applied to axial data, converting these data to directional data by doubling all values *θ*_*i*_ = 2*α*_*i*_. The mean direction was calculated from 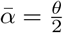, where 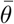 is the common circular mean of the directional data *θ*_*i*_. Similarly, the polarity index (PI) was calculated as the length of the mean resulting vector of directional values *θ*_*i*_. Again, a PI value varies between 0 and 1 and indicates how much the distribution is concentrated around the mean. A value of PI close to 1 implies that the data are concentrated around the mean direction, while a value close to 0 suggests that the data are evenly distributed or random. The axial V-score is computed from

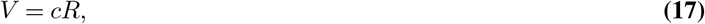

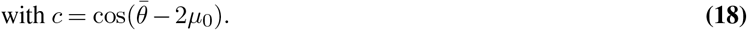

Note, that in this case, the V-score is not the projection of the mean resultant 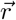 . However, we can compute the following relationship. We assume

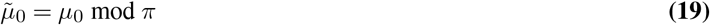

and compute

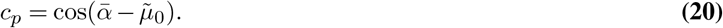

With the commonly known trigonometric identity cos(2*α*) = 2 cos^2^(*α*) *−* 1, we obtain:

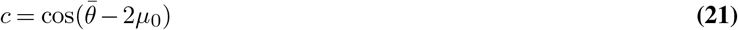

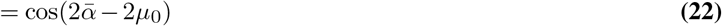

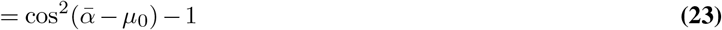

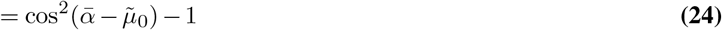

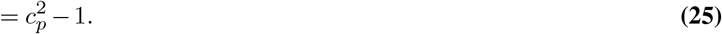

The projection *c*_*p*_ takes values between 0 and 1, with 1 in the case of perfect alignment with the given polar direction and zero for random or perpendicular alignment with the polar direction.

Our web application provides the most common statistical tests for different scenarios, including the Rayleigh test, the V-test, the Rao spacing test and the Watson test applied to directional and axial data (transformed as above). For further discussion of statistical analysis of circular data and extension to comparative statistical analysis of circular data, we refer to (31, 76) and also consider more recent studies (33, 77).

## Supplementary Note 7: Experimental methods and materials

**Supplementary Table S1.** Table of primary and secondary antibodies.

| Antibody name | Host species | Dilution | Product number/manufacturer |
| --- | --- | --- | --- |
| Anti-KLF4 antibody | rabbit | 1:500 | HPA002926 Sigma Aldrich |
| Human VE-Cadherin antibody | goat | 1:1000 | AF938 R&D Systems |
| Anti-GM130 | mouse | 1:500 | 610822 BD Biosciences |
| Notch1 (D1E11) XP® | rabbit | 1:250 | 3608 Cell Signaling |
| Anti-Rabbit-Alexa-647 | donkey | 1:400 | A31573 Invitrogen |
| Anti-Goat-Alexa-568 | donkey | 1:400 | A11057 Invitrogen |
| Anti-Mouse-Alexa-488 | donkey | 1:400 | A21202 Invitrogen |
| Anti-Rabbit-Alexa-488 | donkey | 1:400 | A21206 Invitrogen |

**Supplementary Table S2.**
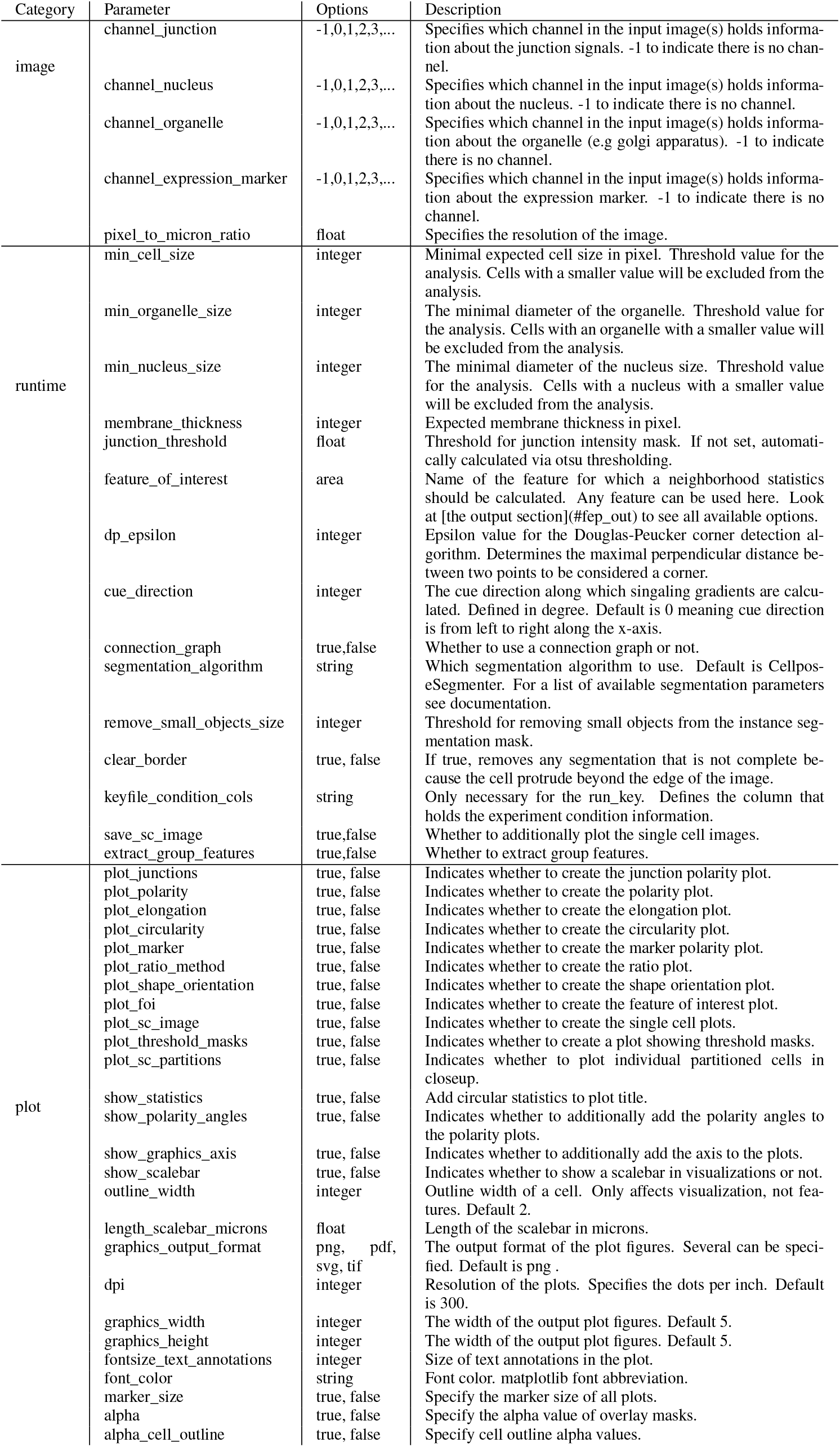
All options and their input specification that can be specified in the parameters file sorted by category.

**Supplementary Table S3.**
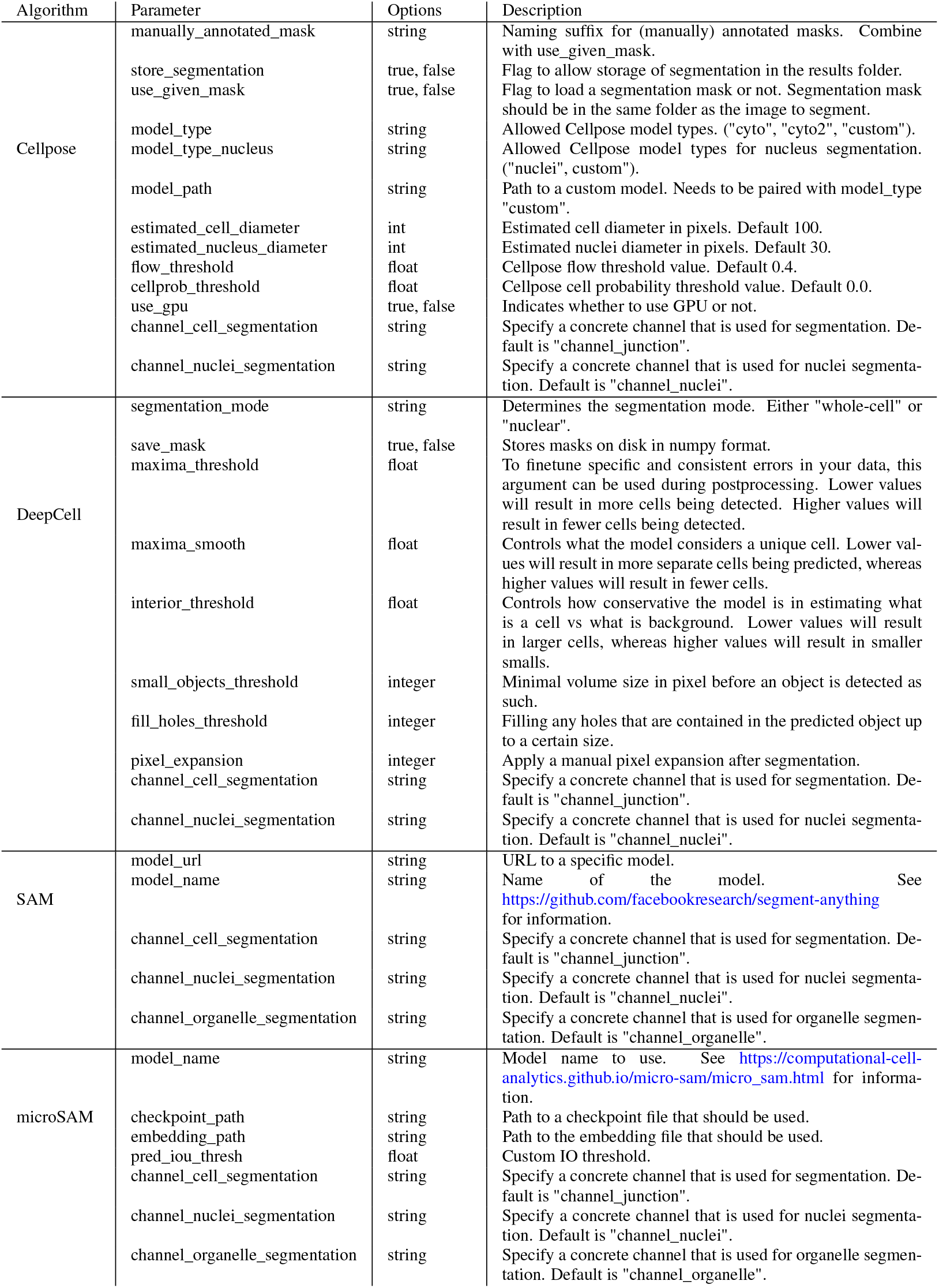
Segmentation parameters for the supported segmentation algorithms. Most parameters are taken from the corresponding publication and/or github repository.

**Supplementary Table S4.**
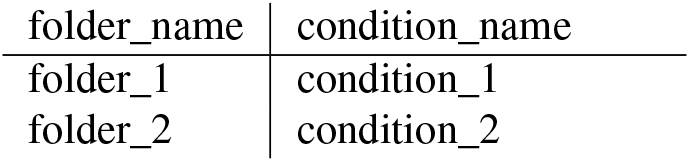
Structure of the *key_file* that can be used to specify experimental conditions and data structure. Note that given folder_names are relative to a root folder passed with the argument named *in_path*.

| folder_name | condition_name |
| --- | --- |
| folder_1 | condition_1 |
| folder_2 | condition_2 |

**Supplementary Table S5.** Categories of features that can be extracted from the image together with their corresponding configuration and explanation.

| Category | Configuration | Explanation |
| --- | --- | --- |
| single cell features | always | The general features extracted from the image. |
| neighborhood features | optional | Statistical properties of a feature of interest (FOI) specified. Includes neighborhood statistics. FOI must be specified. Default is "area". |
| group features | optional | Group properties. Image wise. Includes Morans I correlation analysis. Feature of interest (FOI) must be specified. Default is "area". |
| nucleus features | nucleus channel | All features belonging to the nucleus of the cell. |
| organelle features | organelle channel | All features belonging to the organelle of a cell. |
| junction features | junction channel | All features belonging to the junctions of a cell. |
| marker features | expression marker channel | All features belonging to the expression marker. If a nucleus channel is given, marker nucleus features and marker cytosol features are automatically appended. |

**Supplementary Table S6.**
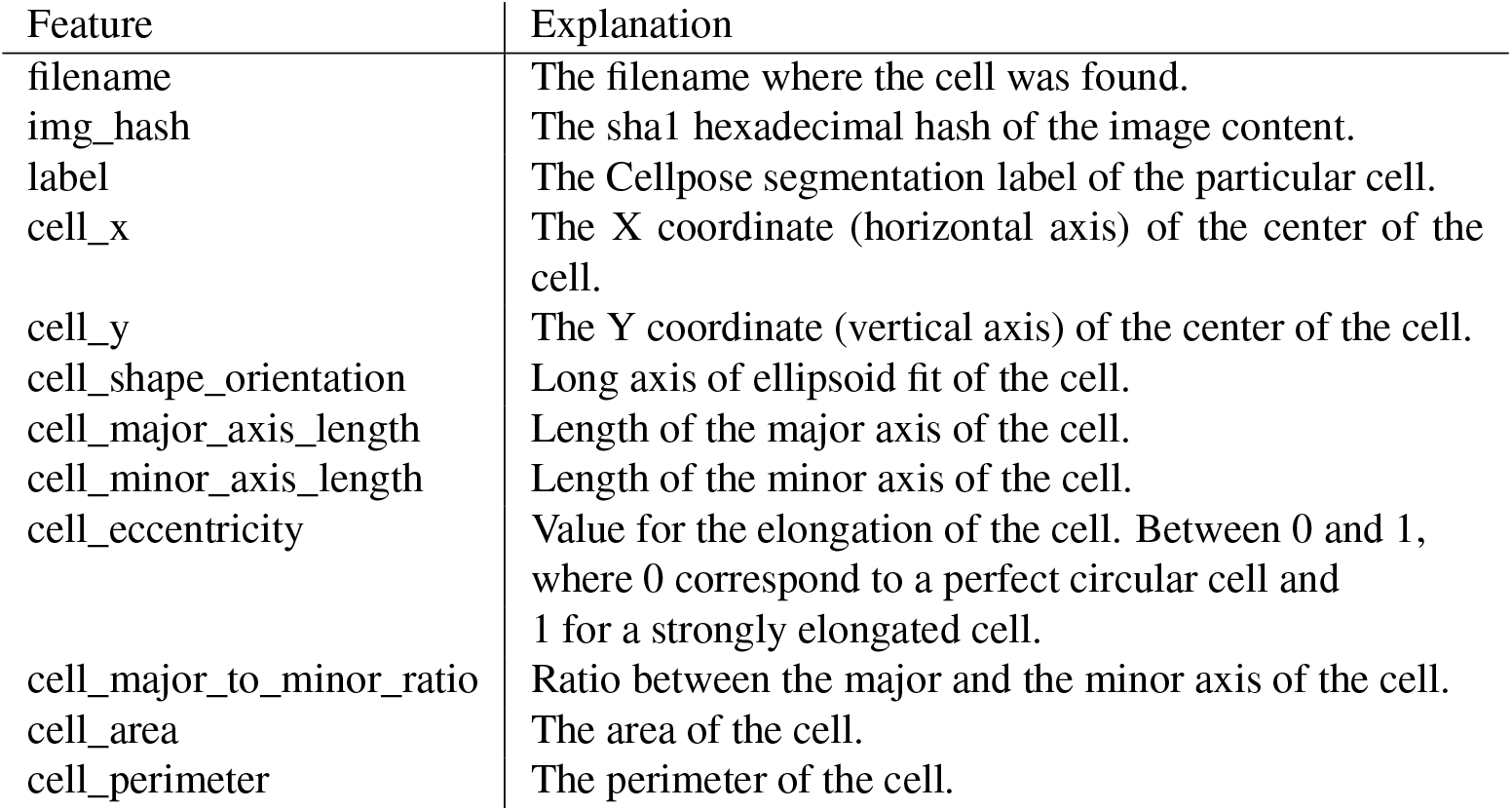
Single cell features features that are extracted from a given image.

**Supplementary Table S7.** Neighborhood features for each cell.

| Feature | Explanation |
| --- | --- |
| neighbors_cell | The absolute number of neighbors of the cell. |
| neighbors_mean_dif_1st | Mean difference of the feature of interest to all first neighbors. |
| neighbors_median_dif_1st | Median difference of the feature of interest to all first neighbors. |
| neighbors_stddev_dif_1st | Standard derivation of the difference of the feature of interest to all first neighbors. |
| neighbors_range_dif_1st | Maximal range of difference of the feature of interest to all first neighbors. |
| neighbors_mean_dif_2nd | Mean difference of the feature of interest to all second neighbors. |
| neighbors_median_dif_2nd | Median difference of the feature of interest to all second neighbors. |
| neighbors_stddev_dif_2nd | Standard derivation of the difference of the feature of interest to all second neighbors. |
| neighbors_range_dif_2nd | Maximal range of difference of the feature of interest to all second neighbors. |

**Supplementary Table S8.** Morans I group statistic performed on a feature of interest (FOI). This is an image wise statistic and is not reported cell wise.

| Feature | Explanation |
| --- | --- |
| morans_i | Statistical correlation analysis |
| morans_p_norm | P-norm of the correlation analysis. |

**Supplementary Table S9.** All nucleus features that can be extracted. Note that a nucleus channel must be configured in the parameters file.

| Feature | Explanation |
| --- | --- |
| nuc_x | X position (horizontal axis) of the cell nucleus. |
| nuc_y | Y position (vertical axis) of the cell nucleus. |
| nuc_disp_orientation_rad | The displacement orientation of the nucleus from the center of the cell in rad |
| nuc_disp_orientation_deg | The displacement orientation of the nucleus from the center of the cell in deg |
| nuc_shape_orientation | Long axis of ellipsoid fit of the nucleus. |
| nuc_major_axis_length | The length of the major axis of the nucleus. |
| nuc_minor_axis_length | The length of the minor axis of the nucleus. |
| nuc_area | The area of the nucleus. |
| nuc_perimeter_nuc | The perimeter of the nucleus. |
| nuc_eccentricity | Value for the elongation of the nucleus. Between 0 and 1, where 0 corresponds to a perfect circular nucleus and 1 to a strongly elongated nucleus. |
| nuc_major_to_minor_ratio | Ratio between the major and the minor axis of the nucleus. |

**Supplementary Table S10.** Organelle features that can be extracted from a given image. Note that an organelle channel must be specified.

| Feature | Explanation |
| --- | --- |
| organelle_x | The X coordinate (horizontal axis) of the center of the cell organelle. |
| organelle_y | The Y coordinate (vertical axis) of the center of the cell organelle. |
| organelle_distance | Distance from cell organelle to the nucleus. |
| organelle_orientation_rad | The orientation in rad of the organelle to the nucleus |
| organelle_orientation_deg | The orientation in deg of the organelle to the nucleus |

**Supplementary Table S11.**
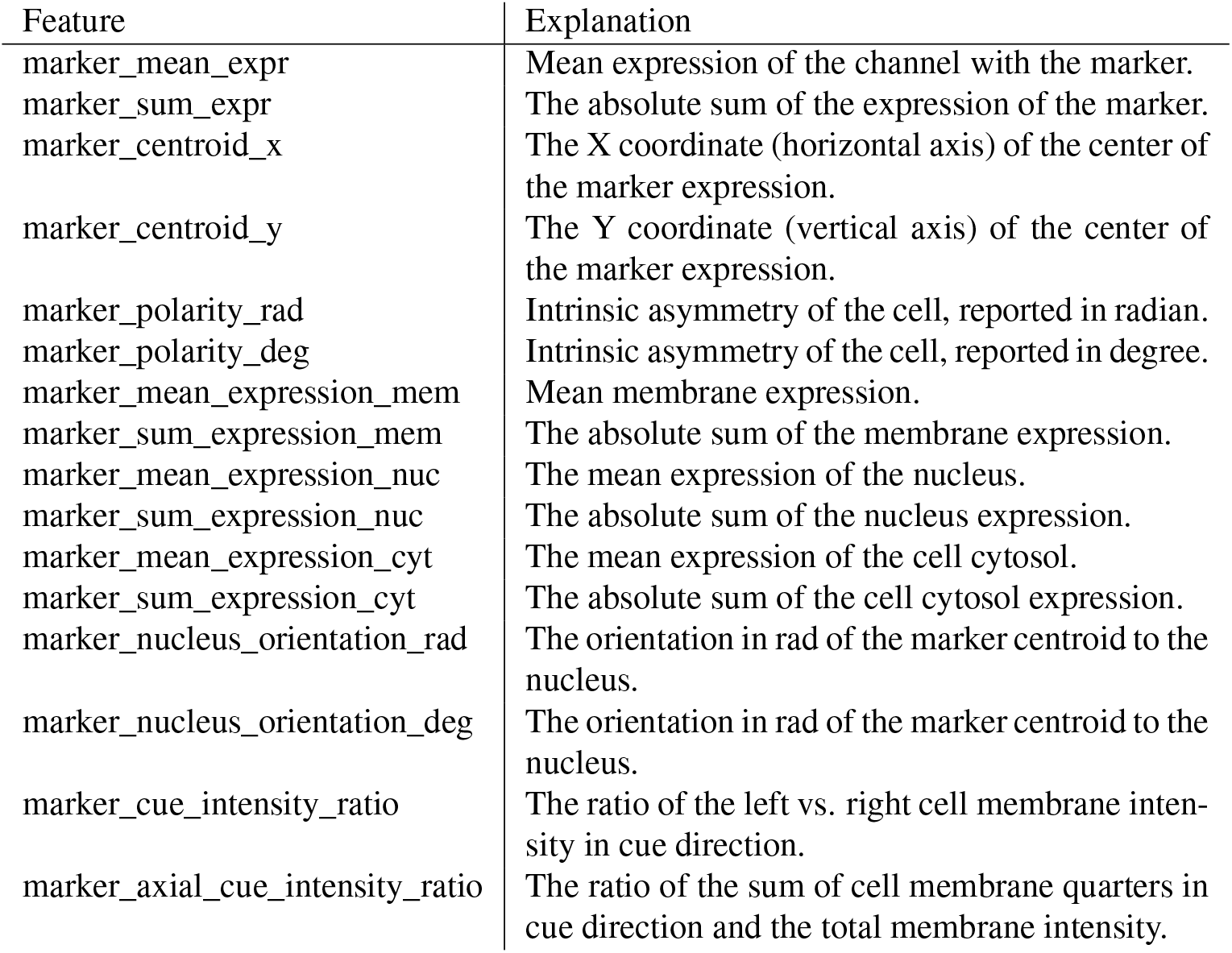
Features that can be gathered if a marker channel is configured.

| Feature | Explanation |
| --- | --- |
| marker_mean_expr | Mean expression of the channel with the marker. |
| marker_sum_expr | The absolute sum of the expression of the marker. |
| marker_centroid_x | The X coordinate (horizontal axis) of the center of the marker expression. |
| marker_centroid_y | The Y coordinate (vertical axis) of the center of the marker expression. |
| marker_polarity_rad | Intrinsic asymmetry of the cell, reported in radian. |
| marker_polarity_deg | Intrinsic asymmetry of the cell, reported in degree. |
| marker_mean_expression_mem | Mean membrane expression. |
| marker_sum_expression_mem | The absolute sum of the membrane expression. |
| marker_mean_expression_nuc | The mean expression of the nucleus. |
| marker_sum_expression_nuc | The absolute sum of the nucleus expression. |
| marker_mean_expression_cyt | The mean expression of the cell cytosol. |
| marker_sum_expression_cyt | The absolute sum of the cell cytosol expression. |
| marker_nucleus_orientation_rad | The orientation in rad of the marker centroid to the nucleus. |
| marker_nucleus_orientation_deg | The orientation in deg of the marker centroid to the nucleus. |
| marker_cue_intensity_ratio | The ratio of the left vs. right cell membrane intensity in cue direction. |
| marker_axial_cue_intensity_ratio | The ratio of the sum of cell membrane quarters in cue direction and the total membrane intensity. |

**Supplementary Table S12.** Features that can be gathered if a junction channel is configured.

| Feature | Explanation |
| --- | --- |
| junction_centroid_x | The X coordinate (horizontal axis) of the centre of the junction expression. |
| junction_centroid_y | The Y coordinate (vertical axis) of the centre of the junction expression. |
| junction_perimeter | The perimeter of the junction area. |
| junction_protein_area | The area with junction protein expression. |
| junction_mean_intensity | The mean junction intensity value. |
| junction_protein_intensity | The mean protein intensity by area. |
| junction_interface_linearity_index | The linearity index of the junction. |
| junction_interface_occupancy | The ratio between junction area and junction protein area. |
| junction_intensity_per_interface_area | The ratio between the junction protein intensity and the junction area. |
| junction_cluster_density | The ratio of junction protein intensity and junction protein area |
| junction_centroid_orientation_rad | The orientation in rad of the junction intensity area centroid to the centre of the cell. |
| junction_centroid_orientation_deg | The orientation in deg of the junction intensity area centroid to the centre of the cell. |
| junction_cue_intensity_ratio | The ratio of the left vs. right cell membrane intensity in cue direction. |
| junction_axial_cue_intensity_ratio | The ratio of the sum of cell membrane quarters in cue direction and the total membrane intensity. |

**Supplementary Table S13.** Comparison of available tools QuantifyPolarity (19), JunctionMapper (20) and Chesnais et al. (14) and our tool.

|  | Polarity-JaM | Quantify-Polarity | Junction-Mapper | Chesnais et al. 2020 |
| --- | --- | --- | --- | --- |
| segmentation | flexible, Cellpose (Deep Learning model) native | specific watershed algorithm (78) | Filter-based | Cell Profiler workflow |
| organelle asymmetries | nuclei-Golgi orientation, polarity index | no | no | no |
| shape asymmetry | size, shape, eccentricity, orientation, topology | size, shape, eccentricity, orientation, topology | no | area, perimeter, shape descriptors and cell neighbours |
| signaling gradients & marker expression | mean intensity in the cytosol, nuclei, and on the cell membrane as well as their ratios | no | no | Notch intensity |
| junction morphology | intensity of junctional markers | intensity of junctional markers | morphology and intensity features | unsupervised classification of junctional morphology of arterial, venous, and microvascular endothelial cell populations |
| polarity measures | 2-axial polarity based on intensity of junctional markers | nuclei-Golgi; polarity of cytosolic and junctional markers, 2-axial cell and nuclei shape orientation | no | no |

**Supplementary Table S14.** Glossary

|  |  |
| --- | --- |
| Deep learning | A family of machine-learning methods, based on deep neural networks (DNN), which are capable of learning representations from data with increasing levels of abstraction (79). |
| Deep Neural Network (DNN) | Computing system inspired by the biological neural networks consisting of multiple (deep) layers between the input and output layers. |
| Instance segmentation | Instance segmentation is a computer vision task that involves identifying and separating individual objects (e.g. biological cells or organelles) within an image, including detecting the boundaries of each object and assigning a unique labels. The result is a pixel-wise map of the image, where each pixel is associated with a specific object instance. |
| Polarity index | Length of the mean resultant vector computed by summing up all single orientation vectors and dividing by the number of measurements. The direction of the mean resultant vector mean angle of the circular distribution and its length the spread. |
| V-score | Similar to the polarity index, but also takes a pre-defined direction into account. |
| Application Programming Interface (API) | Generally, an API describes the connection between different computer programs or tools. It can also refer to everything an application programmer (e.g. python user of Polarity-JaM) needs to know about a piece of code and how to use it. |
| YAML, yaml or yml | A human-readable data serialisation language. Mainly used for configuration files and whenever data need to be stored or transmitted. |

